# A comparative evaluation of CRISPR-Cas9 allele editing systems in *Candida auris*: challenging research in a challenging bug

**DOI:** 10.1101/2025.01.09.632232

**Authors:** Dimitrios Sofras, Hans Carolus, Ana Subotić, Celia Lobo Romero, Craig L. Ennis, Aaron D. Hernday, Clarissa J. Nobile, Jeffrey M. Rybak, Patrick Van Dijck

## Abstract

*Candida auris* is an emergent fungal pathogen of significant interest for molecular research because of its unique nosocomial persistence, high stress tolerance and common multidrug resistance. To investigate the molecular mechanisms of these or other phenotypes, a handful of CRISPR-Cas9 based allele editing tools have been optimized for *C. auris*. Nonetheless, allele editing in this species remains a significant challenge, and different systems have different advantages and disadvantages. In this work, we compare four systems to introduce the genetic elements necessary for the production of Cas9 and the guide RNA molecule in the genome of *C. auris,* replacing the *ENO1*, *LEU2* and *HIS1* loci respectively, while the fourth system makes use of an episomal plasmid. We observed that the editing efficiency of all four systems was significantly different and strain dependent. Alarmingly, we did not detect correct integration of linear CRISPR cassette constructs in integration-based systems, in over 4,900 screened transformants. Still, all transformants, whether correctly edited or not, grew on selective nourseothricin media, suggesting common random ectopic integration of the CRISPR cassette. Although the plasmid-based system showed a low transformation success compared to the other systems, it has the highest editing efficiency with 41.9% correct transformants on average. In an attempt to improve editing efficiencies of integration-based systems by silencing the non-homologous end joining (NHEJ) DNA repair pathway, we deleted two main NHEJ factors, *KU70* and *LIG4*. However, no improved editing or targeting efficiencies were detected in *ku7011, lig411,* or *ku7011/lig4Δ* backgrounds. Our research highlights important challenges in precise genome editing of *C. auris* and sheds light on the advantages and limitations of several methods with the aim to guide scientists in selecting the most appropriate tool for molecular work in this enigmatic fungal pathogen.

**Author summary:** *Candida auris* is a rapidly emerging fungal pathogen that poses serious challenges to global healthcare. Understanding the genetic mechanisms that underlie its nosocomial persistence, virulence, multidrug resistance and other traits is essential for developing new treatments and preventing the spread and burden of *C. auris* infections. However, precise genetic manipulation in *C. auris* has proven difficult due to inefficient genome editing tools. This study compares four different CRISPR-based allele editing systems in *C. auris*, identifying their strengths and limitations. The findings provide crucial insights into selecting the best tools for genetic research in *C. auris*, guiding future efforts to combat this formidable pathogen.

## Introduction

*Candida auris,* is an emergent fungal pathogen that was first described in 2009 [1] and has rapidly become a major concern in global healthcare due to its ability to cause outbreaks of drug resistant invasive infections, particularly in healthcare facilities [2]. *C. auris* has drawn much attention because it displays several unique characteristics for a fungus, such as high rates of resistance to multiple antifungal drugs and disinfectants, high stress tolerance, strong skin colonization potential and exceptional nosocomial transmission capacity, combined with the ability to cause serious, often fatal infections in immunocompromised individuals [3]. *C. auris* emerged globally and near simultaneously from different geographic regions, represented by six phylogenetically distinct clades [4–7]. While the genetic variation between isolates within a clade is minimal, significant genomic differences between clades indicate divergent evolution that began thousands of years ago, with the most recent common ancestor within each clade emerging around 360 years ago and outbreak-causing lineages emerging less than 40 years ago [7]. Interestingly, different clades and strains show different tendencies of virulence, resistance and other phenotypes [7–10].

To study the effects of specific genetic variation in *C. auris*, one needs an allele editing system. Since its discovery in 2012 [11], Clustered regularly interspaced short palindromic repeats (CRISPR) - CRISPR-associated (Cas) gene editing systems allow precise manipulation of DNA in all domains of life. After the successful use of CRISPR-Cas9 gene editing in the fungal model organism *Saccharomyces cerevisiae* in 2013 [12], the technology was optimized for use in the most commonly studied fungal pathogen *Candida albicans* [13, 14], mitigating several biological challenges such as the diploid nature of the genome, the lack of a complete sexual cycle and the unusual codon usage [15–17]. These factors, along with inefficient homologous recombination and a paucity of efficient selectable markers, have historically complicated genome manipulation in fungi [18].

To date, several CRISPR-based allele editing systems have been developed for *Candida* species, each with their own advantages and limitations. While some systems rely on the stable or temporary integration of the CRISPR cassette into the genome, others are transient, recyclable, scarless or make use of a plasmid to express the CRISPR components in the cell. Additionally, *in vitro* assembled Cas9-ribonucleoprotein (RNP) complexes have been used as an alternative for expression-based allele editing systems, although these still require introducing a selectable marker, typically near the locus of interest, which can alter the surrounding genomic architecture. Likewise, several CRISPR editing systems have been developed to manipulate *Candida* genomes beyond allele editing, mainly by temporarily or permanently replacing genes by selectable markers to construct gene knock-out strains. Several comprehensive reviews exist that cover the full diversity, applicability, and use of CRISPR-based systems in *Candida* species [15–20]. Overall, the holy grail of allele editing is a system that has a high editing efficiency and does not leave any trace, such as a selective marker or scar, beyond the aimed allelic edit it was built to introduce. A potential issue in any of these systems is the low efficiency of homologous recombination (HR), on which the correct integration of the donor DNA relies, once Cas9 has produced the double-strand breaks (DSBs). Several *Candida* species, such as *Candida glabrata*, preferentially utilize non-homologous end joining (NHEJ) for the repair of double-strand breaks (DSBs), which competes with HR as an alternative repair pathway [21, 22]. To enhance HR efficiency, key factors of the NHEJ pathway, such as Ku70 and Lig4, have been knocked-out, to improve the CRISPR editing efficiency [23–26].

Here, we compare the efficiency of four different CRISPR-Cas9 gene editing systems in three different clade backgrounds of *C. auris*, using a standardized electroporation protocol. We rely on auxotrophy- and PCR-based screenings to evaluate the editing and targeting efficiency of each system. Furthermore, we assess the impact of deleting *KU70* and *LIG4* on several phenotypes and on the CRISPR editing efficiency of each system. Our findings provide a comprehensive evaluation of all existing CRISPR allele editing tools for *C. auris*, of which one is novel and provide a framework for further optimization of *C. auris* genome editing methodologies.

## Results

### Four different allele-editing systems were evaluated, of which one was newly optimized

In this study, we aimed to introduce a nonsense mutation in the *ADE2* gene to compare CRISPR-Cas9 allele editing efficiencies. The *ADE2* gene encodes for phosphoribosylaminoimidazole carboxylase, an enzyme involved in the *de novo* purine biosynthesis pathway, and has been used as a reliable marker for evaluating gene editing methods because disruption of *ADE2* causes the accumulation of a red pigment, which results in a distinct red colony phenotype [27] (**Figure 1A**). In *C. auris,* no distinct red phenotype was observed for *ade2Δ* colonies on standard YPD agar, but *ade2Δ* strains could be identified by replicating colonies on synthetic medium lacking adenine (**Figure S1**, Supplementary). Thus, after electroporation and plating on YPD+nourseothricin (NTC) to select for transformants, the editing efficiency was first evaluated by replicating the colonies on CSM agar without adenine and with nourseothricin (CSM-ade+NTC). Similarly, auxotrophies for leucine or histidine were identified by colony replicating on CSM medium lacking these nutrients but with nourseothricin (CSM-leu+NTC and CSM-his+NTC), to evaluate targeting efficiency of the *LEU2*- and *HIS1*-integration based CRISPR-Cas9 systems respectively (**Figure 1B**). Using this strategy, we evaluated the efficiency of four different CRISPR-Cas9 systems for genome editing in *Candida auris*. The first system is further referred to as the ‘*ENO1* stable integration’ **(*ENO1-SI*)** system of Vyas *et al.* [13], which was first applied in *C. auris* by Kim *et al.* [28]. As its name infers, the *ENO1-SI* system should allow the stable integration of Cas9 and sgRNA expression cassettes into the genome at the *ENO1* locus, enabling continuous Cas9 expression. The second system we employed is the *LEU2-*targetting temporary integration system (***LEUpOUT***) developed by Nguyen *et al.* [14] and optimized for *C. auris* by Ennis *et al.* [29]. In contrast to the *ENO1-SI* system, the *LEUpOUT* system relies on the temporary integration of the CRISPR-Cas9 cassette into the genome, disrupting the *LEU2* locus, which is reconstituted after the successful removal of the cassette from the genome via homologous recombination. Thirdly, we optimized and employed for the first time in *C. auris,* a *HIS1-* targetting temporary integration system (***HIS-FLP***) based on Nguyen *et al.* [14]. The *HIS-FLP* system is based on the same principles as *LEUpOUT* but targets the *HIS1* locus and allows for marker excision post-editing via *FLP* recombinase leaving an FRT scar (*his1Δ::*FRT). Lastly, we evaluated a plasmid-based system: ‘Episomal Plasmid Induced Cas9’ (***EPIC***), which was optimized by Jeffrey Rybak based on an autonomously replicating sequence from *C. parapsilosis* (*CpARS7*) and used in *C. auris* by Carolus & Sofras *et al.* [30]. The EPIC system comprises an episomal plasmid that enables temporary expression of CRISPR components without genomic integration, and since its maintenance depends on nourseothricin selection, removal of the selective pressure allows the plasmid to be lost after the manipulation is done. The main differences between these systems primarily revolve around whether they involve stable or temporary integration of cassettes or not, and the extent to which they leave behind selectable markers or genomic scars.

**Figure 1:**
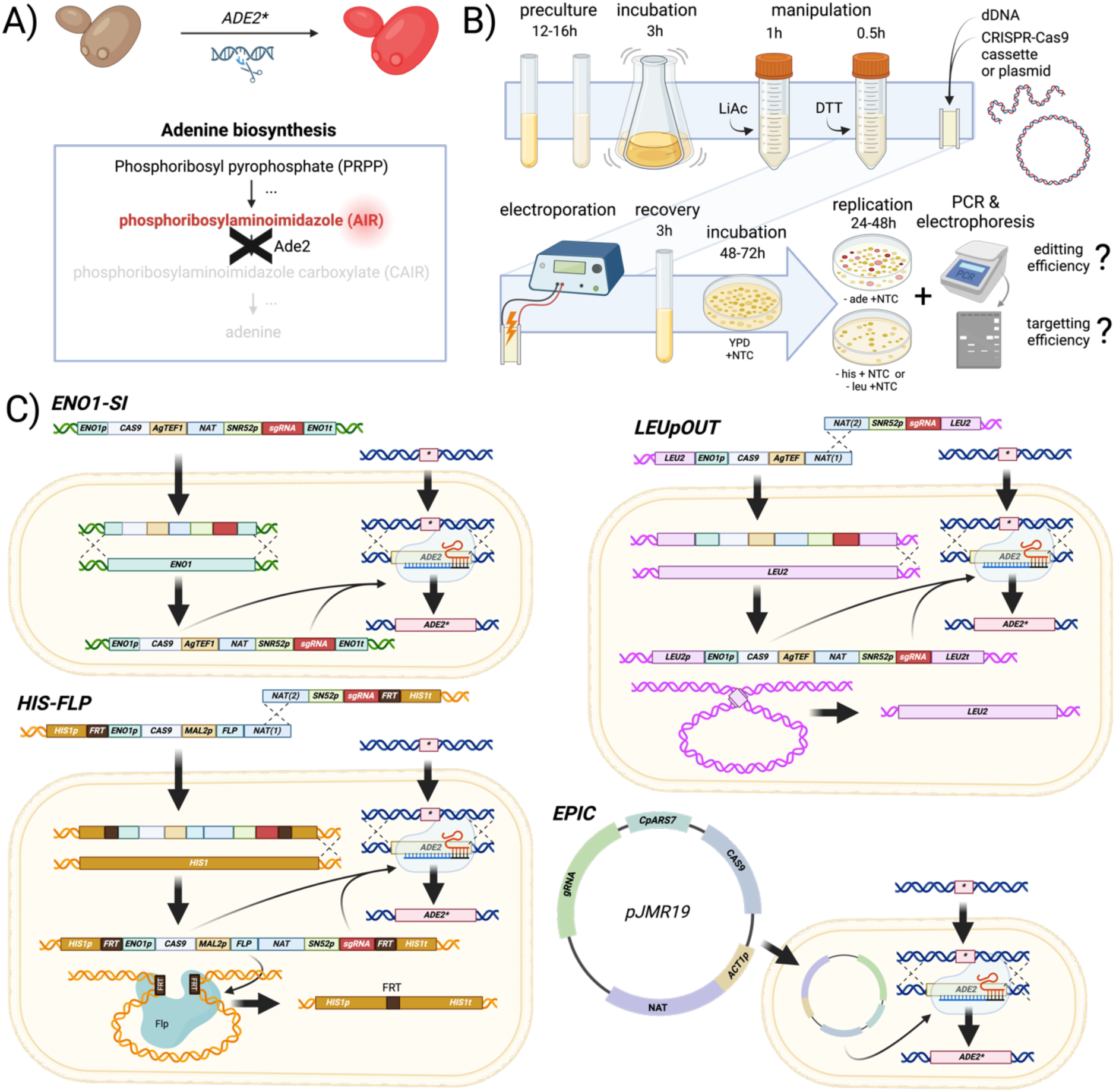
Evaluation of CRISPR-Cas9 systems targeting the *ADE2* gene in *Candida auris*. **(A)** Schematic representation of the *ADE2* gene’s role in the purine biosynthesis pathway, where disruption leads to the accumulation of a red pigment due to a loss of phosphoribosylaminoimidazole carboxylase activity. **(B)** Visual representation of the transformation procedure and evaluation of editing and targeting efficiency. Editing efficiency was evaluated by adenine auxotrophy and verified by PCR for all systems using allele-specific PCR (AS-PCR). Targeting efficiency was evaluated by pooled PCR for *ENO1-SI* transformants and by histidine or leucine auxotrophy for the *HIS-FLP* and *LEUpOUT* transformants respectively. Auxotrophic transformants were PCR verified for all systems. For details, see the *Methods* section. **(C)** Summary of the four CRISPR-Cas9 systems tested for *ADE2* targeting: *ENO1* stable integration (*ENO1-SI*) system, *LEUpOUT* temporary integration system, *HIS1*-targeting temporary integration system (*HIS-FLP*), and the episomal plasmid-based system (EPIC).

All four systems are schematically depicted in **Figure 1C**. A detailed description of all genetic elements of each of these systems can be found in the *Methods* section. As mentioned above, we optimized the *HIS-FLP* system for *C. auris* in this study. In short, we replaced the homology arms of *CaHIS1* for the regions upstream and downstream of *CauHIS1*. In contrast to the other systems of this study, where 500 bp homology regions were used, we opted for 1.5kb of upstream and downstream homology for *HIS-FLP* as a possible solution against ectopic integration (see further). Additionally, the *CaSNR52* promoter sequence was replaced with the respective sequence from *C. auris* to allow for the correct transcription of the gRNA.

### Allele editing efficiency is highly strain- and system-dependent

Due to the importance of strain diversity [31], we included strains from diverse clades of *C. auris*: two strains from Clade I (strain B8441), one of Clade III (strain B11223) and one of Clade IV (strain C52710-20). These clades have shown to be most clinically relevant, causing most invasive and drug-resistant infections [7, 8]. We used two Clade I strains, both originally assigned B8441 (AR0387), but later discovered to show significant differences in certain phenotypes (see further). In one background, used in the Van Dijck lab and referred to as wt I.1, *ku7011,* and ku70*11/lig411* strains were constructed, while in the other background, used in the Nobile lab and referred to as wt I.2, the *lig4Δ* strain was constructed (see further).

Every strain was transformed three times independently, with each system. **Figure 2** shows the editing efficiency of each system for all transformations in each wt background. The editing efficiency was evaluated by the red *ade2-*deficient auxotrophy firstly and then verified by allele-specific PCR (AS-PCR) amplification. Transformants were visually distinguished from ‘background’ colonies since they replicated on YPD+NTC and grew bigger colonies, examples of this are shown in **Figure S2** (Supplementary). The presence and intensity of background growth was highly system dependent with the *ENO1-SI, LEUpOUT* and *HIS-FLP* systems showing a lot of background growth, while background growth was absent in the EPIC system. Also, differences between different strains were observed, with the strain from Clade III exhibiting the highest amount of background growth, followed by wt I.1, wt I.2 and lastly the strain from Clade IV.

**Figure 2:**
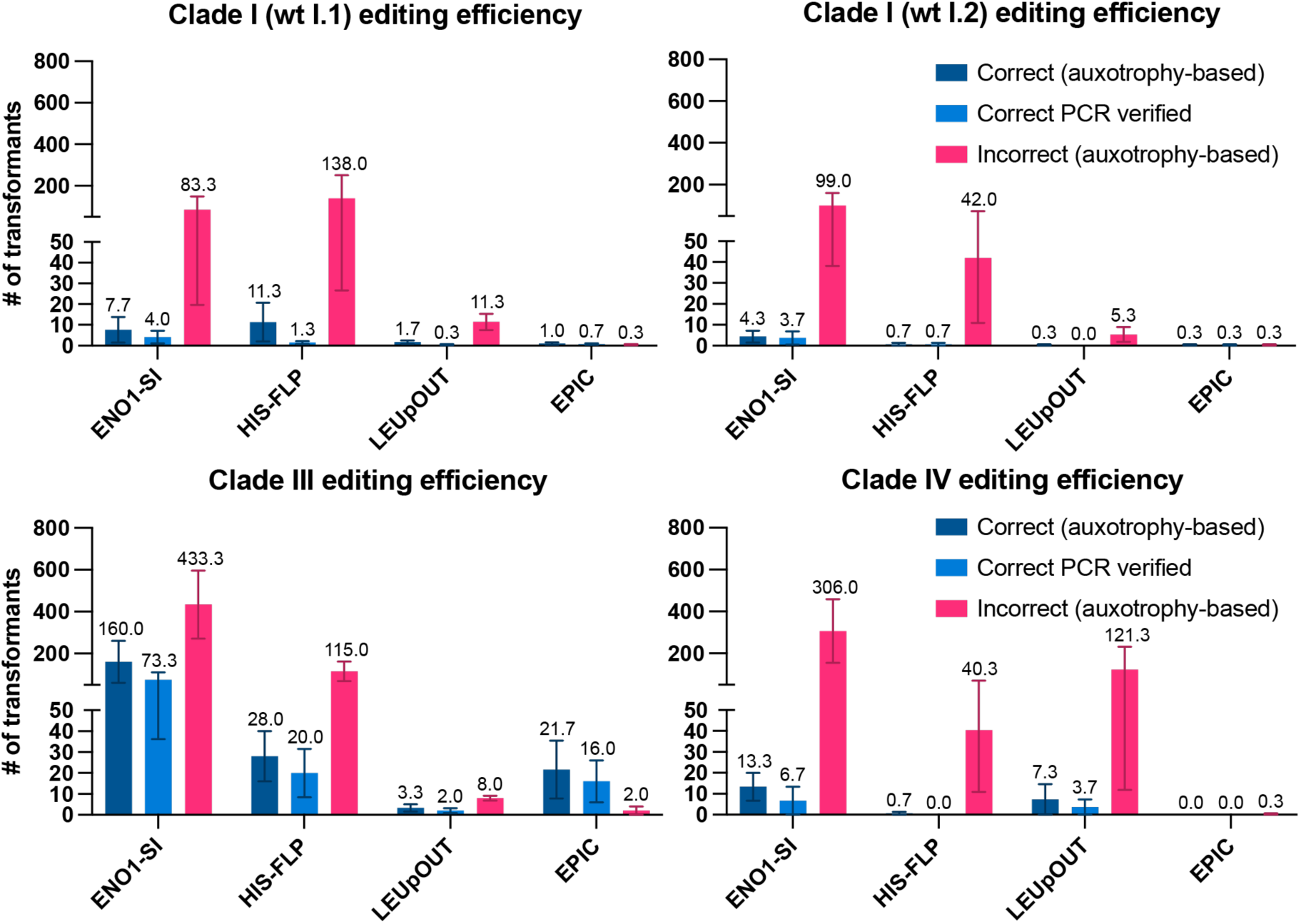
Editing efficiencies of CRISPR-Cas9 systems. Bar graphs depict the editing accuracy of the introduction of a stop codon in the *ADE2* locus using each of the four CRISPR-Cas9 systems, represented by three metrics: correct transformants based on auxotrophy (inability to grow on CSM-ade+NTC medium), correct transformants verified by allele-specific PCR, and the number of incorrect transformants (transformants that grow on CSM-ade+NTC). The efficiencies are presented in four graphs, one for each wild-type (wt) strain used. Each transformation was performed in triplicate, with the mean number of transformants shown on top of the bars. Error bars represent the standard error of the mean (SEM). Photographs of gel electrophoresis runs of all PCRs are shown in **Figure S3** (Supplementary). The source data of this figure can be found in **Table S2** (Supplementary).

The overall number of transformants per transformation round for all strains was lowest for the EPIC system with 6.5 transformants on average, followed by *LEUpOUT, HIS-FLP* and *ENO1-SI* with an average of 39.7, 94 and 276.8 transformants among the four strains tested. The percentage of PCR verified correct transformants was however highest for EPIC, with 41.9% being correct on average, followed by *LEUpOUT* (5.8%), *ENO1-SI* (5.6%) and *HIS-FLP* (4.1%). Overall, the PCR-verified editing efficiency is remarkably low for all cassette-based systems in each background, ranging from 0% (*LEUpOUT* in wt I.2 and *HIS-FLP* in wt of Clade IV) to 17.7% (*LEUpOUT* in wt of Clade III), while the plasmid-based EPIC system had the highest efficiency, with 50% or more correct transformants in Clade I and III strains, although this system did not yield any correct transformants in the Clade IV wt background. Both the number of transformants and the editing efficiency was highest in the Clade III wt background, with 27.9% correct editing in 578.5 transformants on average for all systems, compared to other strains. In the Clade IV background, editing efficiency was the lowest, with 1.2% correct transformants in 367 transformants on average, followed by 13.8 % and 14.5% correct transformants in an average of 114.3 and 191 transformants in wt I.2 and wt I.1 strains respectively. Overall, both Clade I wt strains showed similar editing efficiency results. The PCR verification showed that from all systems and in all strains, 14.3% of all transformants were correct on average. This is remarkably lower than the 21.5% correct transformants based on auxotrophy screening, suggesting that off-target *ADE2*-disruptive edits occurred in 7.2% of the auxotrophic transformants.

Since the transformation success and editing efficiency were background-dependent, we evaluated whether the different background strains showed a difference in survival during the transformation procedure. We estimated the total number of surviving cells by plating on YPD agar and CFU enumeration after initial incubation, adding LiAc, adding DTT, electroporation and recovery, but did not detect a major difference in surviving population sizes between the different strains from the different clades (**Figure S4**, Supplementary). The Clade I wt I.2 strain was not included in this analysis. The differences in total and correct transformants observed in **Figure 2** are thus not related to differential susceptibility of the strains to the transformation procedure and could be potentially due to differences in the cellular uptake, incorporation and/or expression of the CRISPR-Cas9 elements, DNA DSB repair, NTC susceptibility or other aspects.

### Accurate cassette integration is highly challenging

Next, we evaluated the rate of correct cassette integration for the cassette-integration based allele editing systems *ENO1-SI, HIS-FLP* and *LEUpOUT.* For *HIS-FLP* and *LEUpOUT,* transformants were first screened by replica plating directly from YPD+NTC transformation plates onto minimal selective medium with NTC lacking histidine or leucine respectively. Auxotrophic colonies were further tested by colony PCR to assess integration of the CRISPR cassette. For *ENO1-SI,* DNA from all transformants per transformation plate was pooled and PCR-verified right away. To prevent conclusions from false negative PCRs, we targeted the region of interest with 4 different primer pairs as described in the *Methods* section (also see **Table S1** and **Figure S8**, Supplementary), From auxotrophic plate replication, an average of 7.7% and 0.3% of *LEUpOUT* transformants appeared to be leucine auxotrophs in the Clade I (wt I.1) and Clade IV backgrounds respectively, while 2.3% of *HIS-FLP* transformants in Clade III appeared to be histidine auxotrophs. After verification by PCR however, these transformants did not yield the expected bands for targeted integration of the cassettes as shown in **Figure 3**. In addition to the colonies initially identified as auxotrophic transformants, we PCR verified all correctly edited transformants from **Figure 2** for the *LEUpOUT* and *HIS-FLP* transformations, but none showed correct targeting (**Figure S5**, Supplementary). Of the two leucine auxotrophic transformants, none were adenine auxotrophs or showed correct cPCR results, suggesting that the allelic edit of interest did not take place in this subset of transformants. Overall, we were unable to generate positive PCR products for integration of the CRISPR cassettes at the *HIS1*, *LEU2*, or *ENO1* loci, and observed positive bands for the native target loci in all of the tested transformants, except for two *ADE2* wt colonies that were auxotrophs for leucine and showed a band for the integration of the cassette into the *LEU2* locus, but only at the downstream junction. We note that due to the direct repeats that would be generated by integration of the *LEUpOUT* cassette at the *LEU2* locus, it is possible that our PCR primers for the native *LEU2* locus could not yield a false positive result in strains that had the correctly targeted integration, as their length would not allow for their PCR amplification with the set extension time.

**Figure 3:**
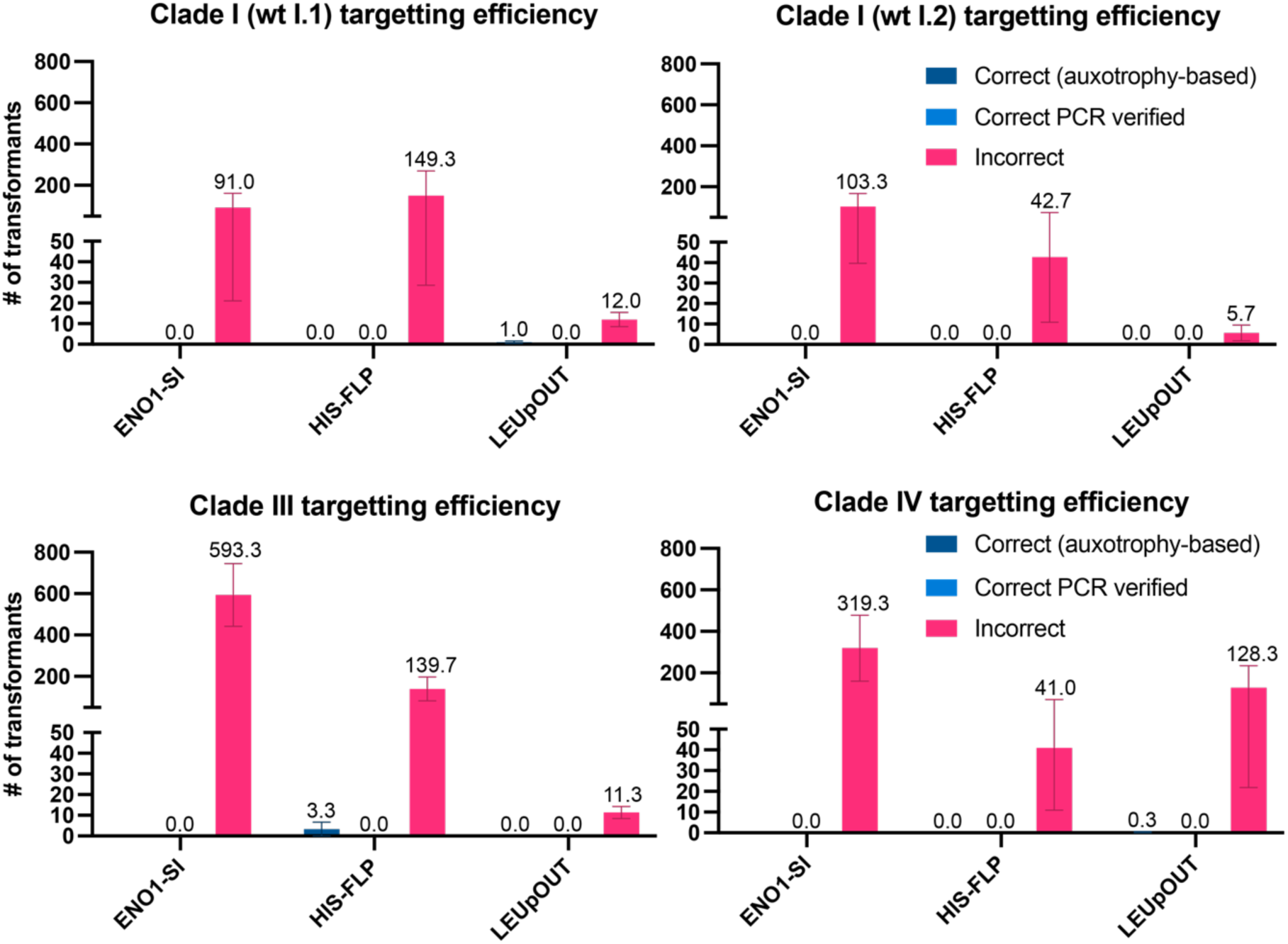
Targeting efficiencies of CRISPR-Cas9 systems. Bar graphs depict the targeting accuracy of the systems that rely on genomic integration: *ENO1-SI*, *HIS-FLP* and *LEUpOUT* in 4 wt strains. For the systems that lead to an auxotrophic phenotype (*HIS-FLP* and *LEUpOUT*), the first screening for correct integration was done by testing the inability of the transformants to grow on CSM-his+NTC and CSM-leu+NTC medium respectively. Final confirmation of correct genomic integration for all systems was verified by PCR analysis. For *ENO1-SI*, DNA of transformants was pooled and screened by PCR, as auxotrophic selection was not applicable. Each transformation was performed in triplicate, with the mean number of transformants shown for each metric on top of the bars. Error bars represent the standard error of the mean (SEM). Photographs of gel electrophoresis runs of all PCRs are shown in **Figure S5** (Supplementary). The source data of this figure can be found in **Table S2** (Supplementary).

Interestingly, all false positive transformants (i.e. NTC resistant transformants that did not contain the cassette in the correct locus and carried either an *ADE2* mutant or wt allele, excluding micro-colonies which were considered true background growth, see Supplementary **Figure S2**) that were checked for the presence of the *NAT* marker by PCR (using primers targeting the *NAT* gene), did show a band of the correct size and thus contain at least one copy of the cassette or *NAT* gene (data not shown). This suggests a systematic failure of our transformants to undergo correct integration of the CRISPR cassette, while the cassette integrates ectopically to maintain NTC resistance during both successful and unsuccessful CRISPR-Cas9 allele editing. We did not further investigate the integrity, copy number, or site of ectopic integration of these cassettes. Since we did not detect any correctly targeted transformants in the *LEUpOUT* and *HIS-FLP* systems, we did not attempt to evaluate the ability or efficiency of recycling these cassettes as demonstrated in literature [14, 29].

Due to the high level of background growth on our transformation plates, untransformed wild-type cells growing in the proximity of NTC resistant transformants could potentially be transferred to the dropout media during replication, and thus appear as false negative prototrophic growth if selection by NTC is not strong enough or NTC is broken down by transformant cells. Nevertheless, such colonies would give ambiguous cPCR results, which were not observed. Another potential hurdle is that selection for both NTC resistance and leucine prototrophy during plate replication, could select for transformants that have spontaneously reconstituted the *LEU2* ORF while simultaneously retaining the NTC resistance marker, either through the generation of mixed-genotype colonies or unintended aneuploidy events. For the *HIS-FLP* system, this is however not possible, as correct integration of the CRISPR cassette results in deletion of the *HIS1* ORF, and spontaneous recycling would result in a scar rendering *HIS1* dysfunctional. Since the rate of incorrect targeting for both *HIS-FLP* and *LEUpOUT* is similar, we hypothesize that false negative plate replication plays a minor or no role. At last, we anticipated that *ADE2* mutant transformants may potentially co-transfer with wild-type colonies, leading to the formation of mixed colonies, however mixed colonies of red *ADE2* deficient colonies and white colonies, nor ambiguous AS-PCR results where observed in this study.

### Suppressing NHEJ does not improve editing or targeting efficiency success

Successful CRISPR-Cas9 editing relies on homology-directed repair (HDR) using a donor DNA (dDNA) fragment, to repair the by Cas9 introduced DSB in the locus of interest. However, NHEJ is an alternative repair mechanism that can inhibit HDR and thus decrease the CRISPR-Cas9 editing efficiency. Several studies have shown that the deletion of *KU70* and *LIG4*, important players in NHEJ (**Figure 4A)**, can lead to improved CRISPR editing efficiency, and increased HDR-mediated targeting efficiency of linear constructs in *Candida* sp. [23–26]. This has so far not been demonstrated in *C. auris,* and the role NHEJ plays in DNA damage repair is unknown in this species. We deleted both *KU70* and *LIG4* independently as well as in combination in the Clade I reference strain B8441 to potentially increase the CRISPR-Cas9 editing and/or targeting efficiency. The *lig411* strains were constructed using a hygromycin deletion cassette by the Nobile group, while the *ku7011* and *ku7011/lig411/* strains were constructed using a *SAT1-*flipper cassette by the Van Dijck group, as described in the *Methods* section.

**Figure 4:**
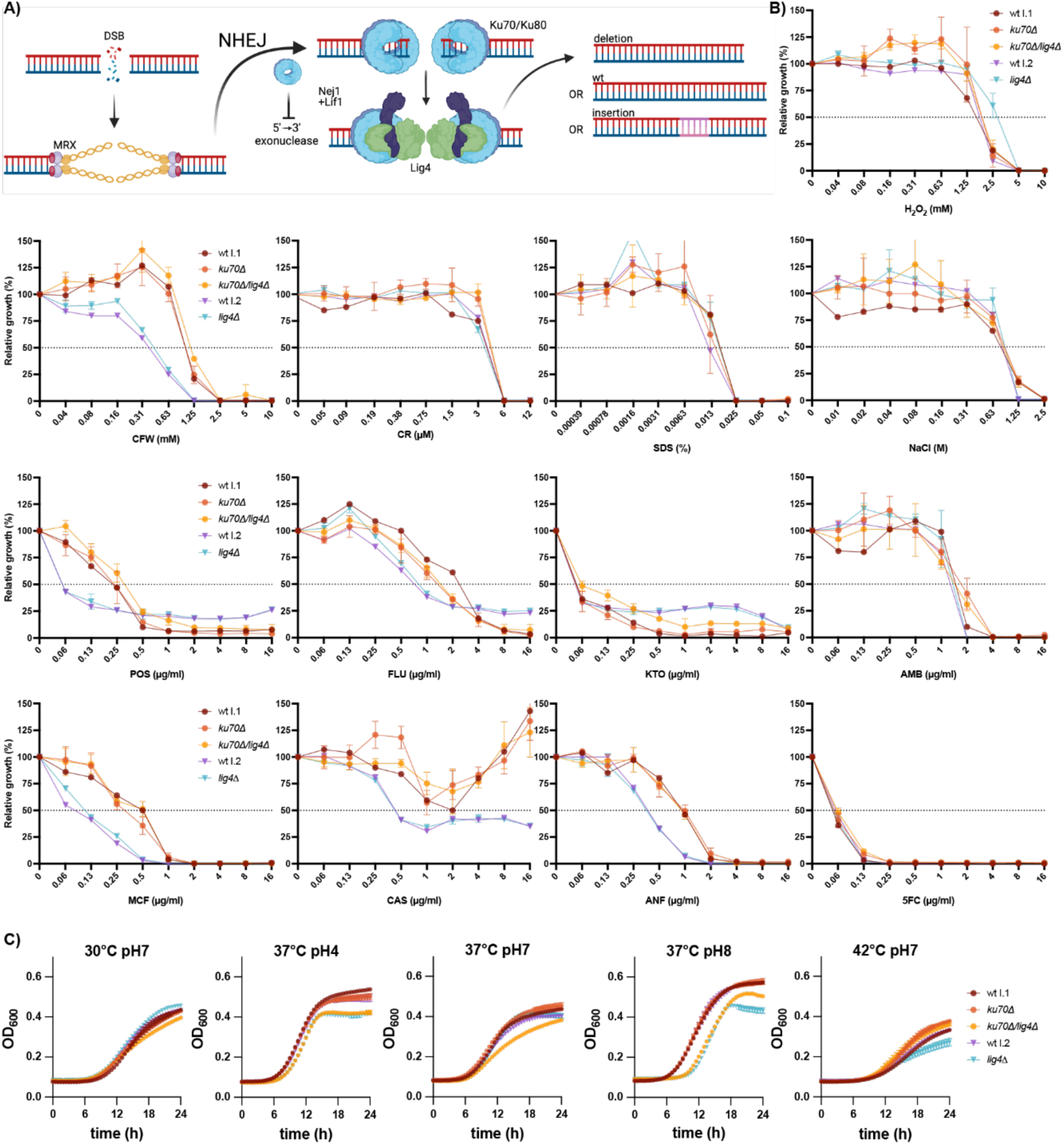
Phenotypic characterization of the *ku70Δ*, *lig4Δ* and *ku70Δ/lig4Δ* strains. **(A)** Non-Homologous End Joining (NHEJ) is one of the two main DNA repair mechanisms. When a double-strand break (DSB) occurs (e.g. due to Cas9 activity), the MRX complex is recruited and plays important roles for both possible repair mechanisms. The first main repair mechanism is homologous recombination (HR) which relies on the initial resection of the DNA strands, which is carried out by a 5’-3’ exonuclease. In NHEJ, the Ku70/Ku80 dimer protects the DNA ends from exonucleases, and is thus necessary for NHEJ. Finally, a protein complex (DNA ligase IV, Lig4) is recruited to ligate the break. Because of the lack of a template, NHEJ often leads to indels of nucleotides. **(B)** Broth dilution assays (BDA) depicted as the relative growth in function of drug/stress concentration of wt I.1 and its derivative strains *ku70Δ* and *ku70Δ/lig4Δ*, and wt I.2 and its derivative *lig4Δ* in RPMI-MOPS (pH 7, 2% glucose) after 48h of incubation. The drugs used were posaconazole (POS), fluconazole (FLU), ketoconazole (KTO), amphotericin B (AMB), micafungin (MCF), caspofungin (CAS), anidulafungin (ANF) and 5-fluorocytosine (5FC). The stress inducing compounds used were calcofluor white (CFW), Congo red, (CR), sodium dodecyl sulfate (SDS), sodium chloride (NaCl) and hydrogen peroxide (H2O2). Each datapoint represents the mean of all biological (n=1 for wt I.1 and wt I.2, n=2 for *lig4Δ*, and n=3 for *ku70Δ* and *ku70Δ/lig4Δ*) and technical (n=2) repeats. Error bars represent the standard deviation (SD). **Figure S6** (Supplementary), shows ETEST results for AMB, CAS, 5FC, and FLU for all strains. **(C)** Growth curves of wt and deletion strains. Growth was measured in RPMI medium with 0.2% glucose under the following conditions: 30°C, 37°C, and 42°C at pH 7, and 37°C at pH 4 and pH 8. Each data point represents the mean of all biological (n=1 for wt I.1 and wt I.2, n=2 for *lig4Δ*, and n=3 for *ku70Δ* and *ku70Δ/lig4Δ*) and technical (n=3) replicates for each strain. Error bars represent the standard error of the mean (SEM).

Before evaluating the effect on CRISPR efficiency, we investigated whether the deletion of these genes affects stress tolerance, drug susceptibility and growth under various conditions. We evaluated susceptibility to five stressors [cell wall stressors calcofluor white (CFW) and Congo red (CR), membrane stressor sodium dodecyl sulphate (SDS), oxidative stressor hydrogen peroxide (H_2_O_2_) and osmotic stressor sodium chloride (NaCl)] and eight antifungal drugs [posaconazole (POS), fluconazole (FLU), ketoconazole (KTO), amphotericin B (AMB), micafungin (MCF), caspofungin (CAS), anidulafungin (ANF) and 5-fluorocytosine (5FC)], while the growth was evaluated over 24h at three temperatures and at three pH levels. Suppressing NHEJ by gene deletions in order to optimize genome editing tools, like CRISPR-Cas9 allele editing, should not be accompanied with strong phenotypic effects, since they could distort the interpretation of biological effects of the alleles under investigation. Of note, we observed a difference in azole, echinocandin and calcofluor white susceptibility between the wt strain in which *KU70* or *KU70+LIG4* were deleted (wt from the Van Dijck lab), and the wt strain in which only *LIG4* was deleted (wt from the Nobile lab). Therefore, we considered these two wt strains, which were both originally assigned ‘*C. auris* B8441*’,* as two different wt strains and named them wt I.1 and wt I.2 respectively. It is important to compare these mutants to their respective wt strain in the following analysis. **Figure 4B** and **Figure S6** (Supplementary) shows that there is no clear difference in drug or stress susceptibility between the *ku7011, lig411, ku7011/lig411,* and their respective wt strains, except for H_2_O_2_ stress to which the *lig411* showed a slight increased tolerance. In growth curve analyses shown in **Figure 4C**, the *lig411* and *ku7011/lig411* strains showed a slight growth deficiency in the form of a lower carrying capacity and/or a prolonged lag-phase in pH 4 and pH 8 conditions. Furthermore, the *ku7011/lig411* and the *lig411* strains showed a decreased growth rate in 37°C and 42°C respectively. This makes us conclude that in *C. auris,* the disruption of *lig411* has phenotypic consequences, while *ku7011* does not, for the conditions tested.

Next, we tested whether the *ku7011, lig411* and *ku7011/lig411* strains show a difference in editing or targeting efficiency in the cassette-based CRISPR-Cas9 systems *ENO1-SI, HIS-FLP* and *LEUpOUT.* The plasmid-based EPIC system was not included in this comparison, since it has a fairly high editing efficiency. **Figure 5A** shows that the editing and targeting efficiencies of three independent transformations in the *ku7011, lig411* and/or *ku7011/lig411* strains was not significantly altered, except for the higher editing success of *ADE2* in the *ENO1-SI* system for the *ku70Δ* strain (13% correct), compared to wt I.1. (4.4% correct). Important to note though, is the great variation among the three transformation repeats, in which 2 out of 3 transformations did not yield any correct transformants (see **Table S2**, Supplementary). Surprisingly, we observed a reduced editing efficiency in the *lig4Δ* vs wt I.2 and *ku7011/lig411* vs wt I.1 for both the *ENO1-SI* and *HIS-FLP* systems. Therefore, we conclude that the disruption of *KU70* or *LIG4* does not improve the overall editing efficiency of cassette-based CRISPR-Cas9 allele editing systems in *C. auris,* suggesting that the NHEJ pathway does not play an important role in the low transformation success. As shown in **Figure 5B**, correct targeting of the CRISP-Cas9 cassettes to the *ENO1, HIS1* or *LEU2* loci was not observed in any of the backgrounds, like in **Figure 3**, again stressing a problematic ectopic integration of linear cassettes, that is not improved by blocking NHEJ.

**Figure 5:**
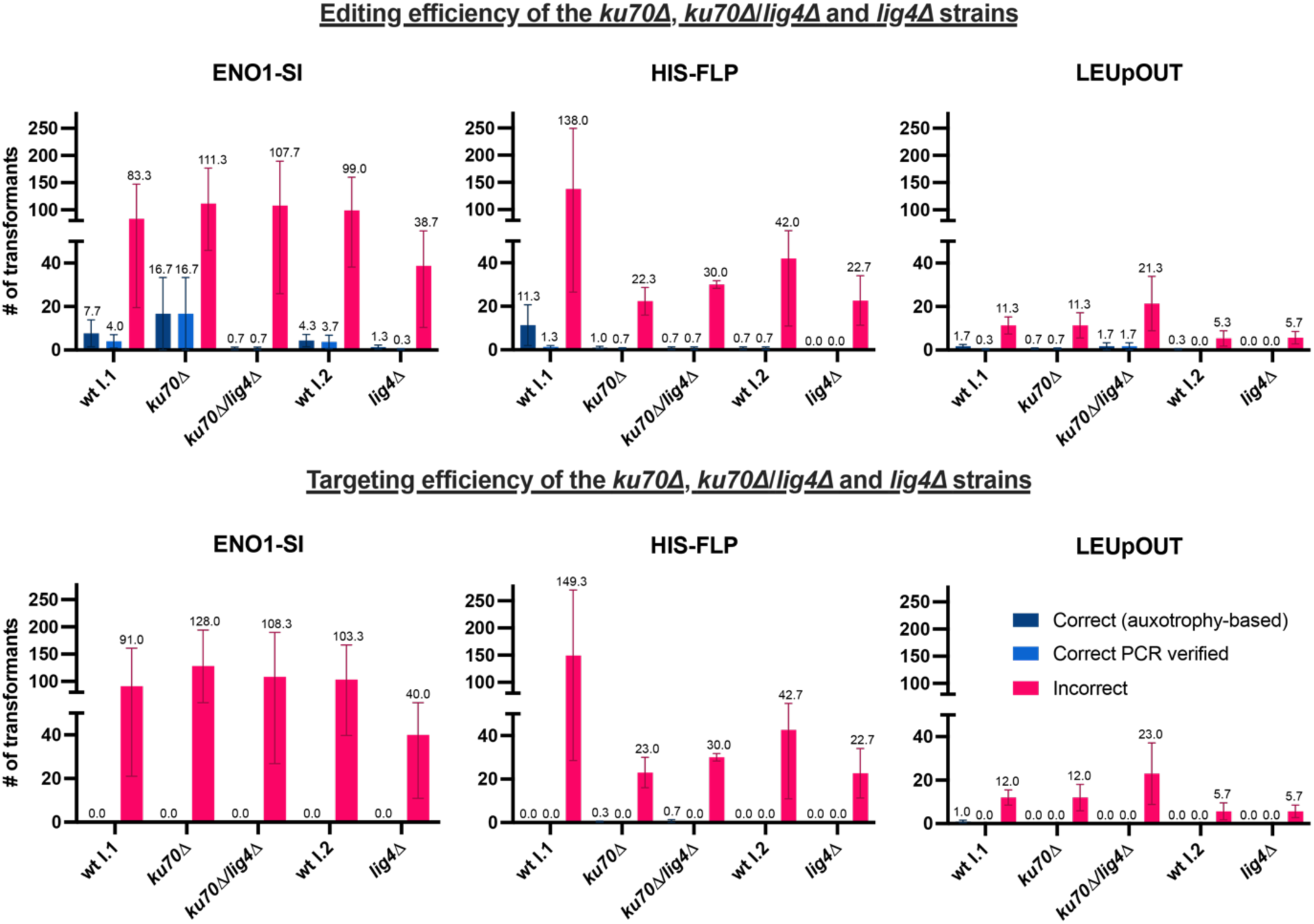
The effect of *ku70* and *lig4* deletions on the editing and targeting efficiencies. **(A)** Bar graphs depict the editing accuracy of the introduction of a stop codon in the *ADE2* locus using the three cassette-integration based CRISPR-Cas9 systems (*ENO1-SI, HIS-FLP* and *LEUpOUT*)-for each genetic background: wt I.1 and its derivative strains *ku7011* and *ku7011/lig411*, and wt I.2 and its derivative strain *lig4Δ*. Editing efficiency is represented by three metrics: correct transformants based on auxotrophy (inability to grow on CSM-ade+NTC medium), correct transformants verified by allele-specific PCR, and the number of incorrect transformants. Each transformation was performed in triplicate, with the mean number of transformants for each metric shown on top of the bar. Error bars represent the standard error of the mean (SEM). **(B)** Targeting accuracy of the same transformations. For *ENO1-SI*, transformants were pooled and screened by PCR, as auxotrophic selection was not applicable. For the systems that lead to an auxotrophic phenotype (HIS-FLP and LEUpOUT), the first screening for correct integration was done by testing the inability of the transformants to grow on CSM-his and CSM-leu, respectively. Final confirmation of correct genomic integration for all systems was verified by PCR analysis. Each transformation was performed in triplicate, with the mean number of transformants shown for each metric. Error bars represent the standard error of the mean (SEM). Photographs of gel electrophoresis runs of all PCRs is shown in **Figure S3** (Supplementary). The source data of this figure can be found in **Table S2** (Supplementary).

## Discussion

In this study, we evaluated four different CRISPR-Cas9 systems for allele editing in *Candida auris*. The *ENO1-SI, LEUpOUT* and *EPIC* systems have been reported before in *C. auris* [28–30], while the *HIS-FLP* system was optimized in this study for *C. auris* based on a strategy used in *C. albicans* [14]. The results of our screening for editing and targeting efficiency in four different *C. auris* strains highlights how challenging genome editing in *C. auris* can be, revealing low but strain- and system-dependent success rates and problematic ectopic cassette integration, underscoring the need for careful system selection and transformant evaluation.

Although the *ENO1-SI* system provided the highest number of transformants, the PCR verified editing efficiency was fairly low (5.6%). The biggest issue with this system was that the cassette was never correctly integrated in the *ENO1* locus. This is potentially due to the essentiality of the *ENO1* gene. *ENO1* encodes for an enolase enzyme with phosphopyruvate hydratase activity, which catalyzes the reversible conversion of 2-phospho-D-glycerate to phosphoenolpyruvate in glycolysis and gluconeogenesis. In *Saccharomyces cerevisiae,* the function of Eno1 can be functionally compensated by its orthologue Eno2, making *ENO1* null mutants viable [32]. In *C. albicans* however, research suggests that only one such enolase gene is present and *ENO1* is essential for growth on glucose [33]. Although *C. albicans ENO1* null mutants are viable on non-fermentable carbon sources, they show reduced drug susceptibility and virulence [34]. In *C. auris,* the essentiality of *ENO1* has not been investigated and no orthologue sequence has been annotated either. This, along with our failed attempts to replace the *ENO1* gene with the CRISPR-Cas9 cassette in 3,321 transformants, suggest that the *ENO1* gene might be essential or important in *C. auris* and thus should not serve as a targeting cassette integration locus. At last, even if the *ENO1* locus would serve as a viable locus for cassette integration, the *ENO1-SI* system is not recyclable, i.e. it is not designed to allow excision of the cassette with selective marker, to remove the expression of the elements encoded on the cassette and introduce another mutation. Thus, even if the cassette integration would work, such system is undesirable.

Both the *LEUpOUT* and *HIS-FLP* systems, which are designed for marker recycling via autonomous and Flp recombinase-mediated recombination respectively, have the major advantage of enabling multiple mutations to be made in consecutive transformation rounds. Theoretically, the *LEUpOUT* system is more desirable than the *HIS-FLP* system, as it is scarless, while the *HIS-FLP* leaves a FRT scar and renders the transformant auxotrophic for histidine. Nevertheless, the *HIS1* locus can be restored using a consecutive transformation round. In our analysis, the *LEUpOUT* system yielded a lower number of transformants compared to the *HIS-FLP* system, although the editing efficiency was higher in *LEUpOUT*. This suggests *LEUpOUT* is superior to *HIS-FLP,* both in terms of design and success rate. Nevertheless, like the *ENO1-SI* system, incorrect integration of the CRISPR cassettes was also a major problem with both the *LEUpOUT* and *HIS-FLP* systems. This highlights that under the transformation conditions that we used, random integration, rather than targeted integration via homologous recombination, appears to prevail, even if presumably non-essential genes such as *LEU2* or *HIS1* are targeted for cassette integration. We note that Ennis *et al.* [29] did not report the frequency of correct integration of the LEUpOUT CRISPR cassette. Furthermore, Ennis et al. observed an across-clade average efficiency of 40% and 99%, respectively, for deleting and restoring the *CAS5* gene at the native locus, indicating that integration of linear DNA fragments at the CRISPR target locus via homology directed repair can occur with high frequencies in *C. auris*. We also note that Ennis et al. observed these higher frequencies of editing and targeting success with the hygromycin-resistant *LEUpOUT* system, as opposed to the nourseothricin-resistant version that we tested (personal communications), suggesting that the combination of strain, system, marker, and transformation protocol may all be critical to successful genome editing via homologous recombination in *C. auris*.

We did identify leucine auxotrophic transformants in this study at extremely low frequencies, but a fully correct recombination of the cassette was never identified by means of various PCRs. This suggests that although recombination occurs, this process is highly error prone under the conditions we tested. This leads to the important question whether critical methodological variables have a much higher influence on the relative frequencies of intended integrations via homologous recombination vs ectopic integration via non-homologous end joining in *C. auris,* as opposed to other *Candida* species where random integration has not been reported to this extent. Many recent studies construct mutants based on homologous recombination of linear cassettes, with or without the help of Cas9-RNP complexes [28, 30, 35–48]. Santana *et al.* used two transient expression cassettes to perform CRISPR in *C. auris* with reportedly high success rates, but we did not include their system in this study due to the obligatory introduction of a selection marker in the vicinity of the desired genetic alteration [49]. However, few studies report adequate controls for correct integration and rarely, problematic transformations are mentioned. Pelletier *et al.* [47] mention how a clean *ALS4112* null mutant could not be obtained. They used an inverse PCR strategy to show that a 21.3kb region, containing six additional genes downstream of their gene of interest, was deleted in the transformant they continued with. Mayr *et al.* [38] report random, ectopic and multicopy integration of the *SAT1* gene deletion cassette in their efforts to construct *MRR1a/b/c* and *TAC1a/b* null mutants in *C. auris,* however they also report that using lower levels of NTC in their transformation plates (50µg/ml vs the typically used 200ug/ml) reduced the ectopic integration issues when modifying these loci. In a pilot study, we also used lower concentrations of NTC, but this did not increase the targeting efficiency, while it led to more background growth on the transformation plate (data not shown). Mayr *et al.* discovered the multicopy integrations by restriction digestion and southern blot analysis [38], which provides a more detailed confirmation of correct integration and can identify more complex genomic changes compared to our PCR-based approach. Nevertheless, PCR-verification of transformants is faster, easier and cheaper for screening transformants, while other methods like blotting or whole genome sequencing are unfit to systematically screen thousands of transformants. Besides low expression of the selective marker (and thus low resistance to the selective agent, like nourseothricin), the length of the homology arms (being too short) has been put forward as potential reasons for the low transformation success and ectopic, multicopy integration [38, 50], although data is lacking to prove this.

We note that in contrast to the Ennis et al LEUpOUT protocol, which relies on chemical transformation using lithium acetate and heat-shock, our study used an electroporation-based transformation protocol which includes multiple washes with ice-cold buffers that is similar to the protocols used by other groups that also observed high levels of ectopic integration in *C. auris* [38, 39]. This raises the possibility that cold stress, or other aspects of these commonly used electroporation protocols, could be driving an increase in random DNA damage in *C. auris*, relative to heat-shock transformation protocols, and thus increasing the frequency of random ectopic integration of linear DNA fragments. We did not assess whether the EPIC plasmid became integrated into the genome or remained episomal under our transformation conditions. However, allelic variants made with *EPIC* in Carolus *et al.* were easily recycled by growing the transformants on non-selective media [30], thus indicating that losing the plasmid is feasible and integration of the selective marker is uncommon. Moreover, circular DNA is less recombinogenic compared to linear DNA [51] and the cassette based systems (*ENO1-SI, HIS-FLP* and *LEUpOUT*) contain larger fragments of endogenous DNA compared to the plasmid based (*EPIC*) system, making genomic integration less likely for *EPIC*.

In summary, any genetic manipulation in *C. auris,* and potentially in other fungal species, which relies on homologous recombination of linear DNA fragments, should be meticulously verified. One cannot simply rely on auxotrophies and single PCR verification (e.g. using only primers within the cassette or within the gene), which are prone to false positive and false negative results, to verify recombination-based edits. Instead, one should resort to multiple PCRs, targeting the amplification of regions spanning the gene, cassette, upstream and downstream region (up/downstream of the homology region of the cassette/target) as conducted here, or restriction and southern blot analysis as reported by Mayr *et al.* [38]. Ultimately, one should always sequence the targeted region, as NHEJ and recombination of dDNAs can always lead to off-target modifications that can go undetected by PCR or hybridization-based methods. Alternatively, long read whole genome sequencing methods could verify correct genetic edits and ectopic integrations. Given this complexity, the use of multiple independently constructed transformants in experiments, and reporting outliers in this effort, is important to consider in any constructed-mutant based research. The use of biological repeats (multiple transformants) mitigates the potential off-target effects ectopic integration can cause more than a strategy in which re-integrant/allele-restoration transformants are used to investigate gene or allele functions.

Given the low frequency of targeted genome editing and the high frequency of incorrect cassette integration that we observed with the cassette-based systems, *EPIC* [30] emerges as the most reliable choice for *C. auris* allele editing under the electroporation-based transformation conditions that we tested. The episomal nature of *EPIC* may reduce the risk of ectopic, error-prone integration of linear DNA fragments, providing a safer and more efficient alternative for researchers aiming for precise genome manipulations. EPIC has the added value of introducing only one selective marker in the cell, without disrupting genes that are essential in certain (nutrient-lacking) conditions, which could otherwise impose additional stress on recovering transformed cells. The number of transformants using *EPIC* was however significantly lower compared to the cassette-based systems, limiting its overall throughput. Attempts for potential improvements such as the use of protoplasts to improve plasmid uptake, the use of alternative transformation procedures such as heat shock-based methods, or further optimization of the plasmid, can only be encouraged. Regardless of the low number of transformants, the *EPIC* system showed the highest relative editing efficiency, with 40% or more of the transformants being correctly edited, although no transformants were obtained for the Clade IV wt strain. The latter showcases the strain-dependent variation in transformation success and CRISPR efficiency, which has been reported before in *C. auris* [29, 38]. It is worth mentioning that *EPIC* has been successfully used for allele editing successfully in the same Clade IV wt strain used in Carolus & Sofras *et al.* [30]. In this study, the Clade III wt strain showed the highest transformation success and the Clade IV wt strain showed the lowest transformation success. In Ennis *et al.* [29], a Clade III strain also showed the highest transformation success compared to strains from Clade I, II, IV and V for deleting one gene, although in Mayr *et al.* [38], no significant difference in transformation success between a Clade III and Clade IV strain was reported in deleting several genes. This suggests that the differential transformation success is strain-or target-rather than clade-specific. The discrepancy in transformation yield was not related to survival during the transformation process in our study, suggesting that inherent differences in the DNA repair pathways or chromatin structure between strains or clades may play a significant role. Previous research indicates that species-dependent variation in CRISPR efficiency may be due to differences in the efficiency of homologous recombination and other DNA repair mechanisms, which vary widely between fungal species [18] and potentially within clades or strains of the same species. Alternatively, the way foreign DNA is taken up and expressed might differ. It is important to note that this study did not seek to replicate previously reported CRISPR editing efficiencies or optimize existing protocols. Instead, the primary objective was to perform a comparative evaluation of CRISPR systems applied in *C. auris*, utilizing a standardized transformation protocol as detailed in the *Methods* section.

Despite our attempts to improve CRISPR efficiency by knocking out key components of the non-homologous end joining (NHEJ) pathway (by deleting *KU70* and *LIG4*), no significant improvements were observed. This was unexpected, as impeding NHEJ has been shown to increase homology-directed repair (HDR) and improve targeted homologous recombination in other *Candida* species [23–26]. Interestingly, the deletion of *LIG4* but not *KU70* showed a minor undesirable phenotype, which contrasts Cen *et al.* [25], who report no effects of *lig4Δ* but an effect on stress and drug tolerance in *ku80Δ* in *C. glabrata.* Potentially, Lig4 and the Ku70/Ku80 complex play a different role in *C. auris,* compared to other species. Beyond *KU70* and *LIG4*, additional factors within DNA repair pathways may contribute to the persistence of ectopic integrations in *C. auris*. Proteins involved in the processing of double-strand breaks, such as those in the MRX complex (Mre11, Rad50, Xrs2) and Sae2 [52], play critical roles in determining the balance between homologous recombination (HR) and alternative repair mechanisms like single-strand annealing (SSA) or microhomology-mediated end joining (MMEJ) [22, 53, 54]. These pathways can compete with homology-directed repair (HDR), potentially leading to unintended integrations even in strains deficient in non-homologous end joining (NHEJ). Furthermore, other components of the homologous recombination machinery, such as Rad51 or its regulators, may influence the fidelity of CRISPR-mediated genome editing, as shown for mammalian cells [55].

In conclusion, our results demonstrate that the episomal plasmid-based CRISPR-Cas9 system *EPIC* is the most reliable allele-editing tool for *C. auris* under the electroporation-based transformation conditions we tested. Although it produces fewer transformants compared to the cassette-based systems, *EPIC* achieves the highest rate of accurate edits and has an intrinsically lower chance of genomic integration by design, making it a valuable system for precise genetic manipulation in *C. auris*. Moving forward, future research should focus on optimizing plasmid-based systems, possibly by increasing transformation efficiency through methods like protoplast generation or refining transformation protocols. Additionally, exploring strategies to further enhance homologous recombination efficiency and optimize plasmid gene expression can be extremely useful. These improvements will advance our ability to manipulate the genome of this challenging fungal pathogen and enable deeper insights into the mechanisms underlying its unique biology.

## Methods

### Strains and growth conditions

The parental strains used in this study were single colony isolates from clinical strains of Pakistan (Clade I), South Africa (Clade III), and Colombia (Clade IV). Strain information is listed in **Table S3** (Supplementary). Strains were stocked at −80°C in 20% glycerol and routinely plated on solid YPD (1% w/v yeast extract, 2% w/v bacteriological peptone, 2% dextrose v/v) agar (2%) at 37°C unless stated otherwise.

### Growth curves

Overnight cultures were prepared in RPMI 1640 (Sigma-Aldrich) with 2% glucose and 165mM morpholinepropanesulfonic acid (MOPS, Sigma-Aldrich) buffered at pH 7 with KOH. The cultures were adjusted to a final cell concentration of 10^6^ cells per 200 µL in each well, based on spectrophotometric measurements. Three growth media were used, comprising RPMI 1640 (0.2% glucose) buffered with MOPS (165mM) and KOH at pH 4, 7 and 8. Growth was monitored at 37°C for all pH conditions and at 30°C and 42°C for pH 7 measuring the optical density at 600 nm (OD_600_) using a Multiskan GO automated plate reader (Thermo Scientific) in flat-bottom 96-well microplates (Greiner) with intermittent (10 min. interval) pulsed (1 min medium strength shaking) shaking and 30-minute interval OD_600_ measurements. Growth curves were generated based on three replicate measurements per biological repeat.

### Compound susceptibility testing

Drug susceptibility was assessed by broth dilution assays (BDA). Briefly, a series of nine twofold dilutions of compound was prepared in a final volume of 200 µl RPMI–MOPS (pH 7, 2% glucose, 1% DMSO) medium. 100–500 cells were seeded in each well of a round-bottom 96-well polystyrene microtitre plate (Greiner). Plates were incubated at 37 °C for 48 h, and growth was assessed spectrophotometrically (OD_600_) using a Synergy H1 microplate reader (BioTek). The BDA curves were constructed in Graphpad prism and relative growth equals to the relative growth in each compound concentration to the growth in the untreated condition. Susceptibility to amphotericin B, fluconazole, 5-fluorocytosine, caspofungin, anidulafungin and micafungin was assessed by ETEST (bioMérieux). In short, cotton swabs saturated with cell suspension adjusted to an OD 0.1 was used to spread the cells on MOPS-buffered (165mM, pH 7) RPMI 1640 (2% glucose) agar plates. The plates were incubated at 37 °C and scans were taken at 24 and 48 h.

### Plasmid contstruction

#### SAT1 flipper

To delete *KU70 and LIG4*, SAT1 flipper cassettes were constructed to target these genes. For each gene, a 500 bp sequence upstream of the ORFs was cloned into the linearized pSFS2 vector after digestion with ApaI and XhoI. The intermediate vectors were then digested with NotI and SacII and a 500 bp region downstream of the ORFs was cloned into the linearized vectors using NEBuilder® HiFi.

#### *LIG4* single deletion

The *lig4Δ* (wt I.2) deletion strain was created by replacing the *LIG4* open reading frame with a PCR-amplified hygromycin marker with Phusion® High-Fidelity DNA Polymerase. Cells were grown to mid-log in YPD, washed with sterile water, and incubated overnight in polyethylene glycol, Lithium Acetate, and TE as described in *Ennis et al.* [29]. Transformed cells were washed twice with YPD and recovered for 4 h at 30°C before plating on YPD plates supplemented with hygromycin at 500 µg/ml. Colonies were screened by two sets of oligos to confirm in-frame insertion of the hygromycin marker.

#### *ENO1-SI* (pV1210)

The approach was identical as described in Kim *et al.* [28]. pV1200 [13] as digested with KpnI and XmaI and the *CauENO1p*, amplified from the B8441 (wt I.1) genome, was inserted to the linearized vector using NEBuilder® HiFi (New England Biolabs). The intermediate vector was digested with NotI and SacI, and the *CauSNR52p*, *CauENO1term*, amplified from the B8441 (wt I.1) genome and the gRNA scaffold sequence amplified from pV1200 were inserted in a single ligation round using NEBuilder® HiFi to produce pV1210. The gRNA sequence for targeting *ADE2* was introduced by digesting pV1210 with BsmBI and ligation of duplexed oligos using NEBuilder® HiFi.

#### HIS-FLP (pADH99Cau and pADH100Cau)

For HIS-FLP, there are two plasmids required. pADH99Cau contains the sequences encoding for Cas9, the Flp site-specific recombinase and the first 150 amino acids of nourseothricin N-acetyl transferase (NAT1/2). pADH99Cau also contains the genomic sequence of B8441 (wt I.1) 1500 bp upstream of *HIS1* (B9J08_005247) used as the homology region for genomic integration and the *CauENO1p* driving the expression of Cas9. pADH100Cau contains the sequences encoding for the last 174 amino acids of NAT with an overlap of 134 amino acids with NAT1/2, the *CauSNR52* promoter sequence followed by the gRNA scaffold sequence and the genomic sequence of B8441 (wt I.1) downstream of *HIS1*.

pADH99 was digested with NcoI and XmaI to remove the *CaENO1p* and the *CaHIS1_US* sequences. A 1.5 kb fragment upstream of *HIS1* and the *CauENO1p* amplified from genomic DNA were inserted in the linearized vector using NEBuilder® HiFi to produce pADH99Cau. pADH100 was digested with BsmI and SapI. The *CauSNR52p* and gRNA scaffold sequences, amplified from pV1210, and a 1.5 kb fragment downstream of *HIS1* amplified from genomic DNA were assembled in the linearized vector using NEBuilder® HiFi.

#### LEUpOUT

The vectors pCE35 and pCE27 were constructed by Ennis *et al*.[29].

#### EPIC (pJMR19)

The vector was constructed by Jeffrey Rybak. The gRNA sequence for targeting *ADE2* was introduced by digesting pJMR19 with SapI and ligating duplexed oligos using T4 DNA ligase (New England Biolabs).

All **oligonucleotides** used in this section and for Sanger sequencing to confirm the successful plasmid construction are listed in **Table S1**.

### Transformation protocol

For all the transformations we used the same electroporation protocol, but we varied the DNA concentrations according to the original publication recommendations [14, 28–30]. Single colonies were inoculated in liquid YPD and grown overnight at 37°C in a shaking incubator. The precultures were diluted in 50 mL YPD in a conical flask to an OD_600_ of 0.4 and grown until the OD_600_ reached a range of 1.6 to 2.2 (approximately 3-4 hours). The cells were collected (5 minutes at 3,273 x g), resuspended in 10 mL of transformation buffer [10 mM Tris-HCl, 1 mM EDTA•Na2 (VWR) and 100 mM LiOAc (Sigma)] and shaken at 37°C, 150 rpm for 1 hour. 250 μL of 1 M DTT (VWR) was added, and the cells were incubated for an additional 30 minutes. Cells were washed twice (5 minutes at 3,273 x g at 4°C), first with 25 mL ice-cold dH_2_O and then with 5 mL ice-cold 1 M sorbitol (Sigma). The supernatant was removed carefully, and the pellet was resuspended in 200 μL of ice-cold 1 M sorbitol. 40 μL of the competent cell suspension were mixed with the transformation mixture and transferred in a 2 mm electroporation cuvette (Pulsestar, Westburg). A single pulse was given at 1.8 kV, 200 Ω, 25 μF, and the transformation mixture was immediately transferred to 2 mL YPD in test tubes following incubation for 4 hours at 37°C, 150 rpm. The cells were collected by centrifugation of 5 minutes at 5,000 x g, resuspended in YPD and plated on YPD agar containing 200 mg/mL of nourseothricin (Jena bioscience) in 1:1, 1:10 and a 1:100 dilutions. Transformants appeared after two to three days of incubation at 37°C.

#### SAT1 flipper

The constructed vectors were linearized by digestion with KpnI and SacII and the deletion cassettes were purified from a 1% agarose gel using the Wizard® PCR and SV Gel Clean-Up System (Promega). 500 ng of the cassette was used in each transformation round. Correct deletion mutants were confirmed by PCRs of the upstream and downstream junctions of the *KU70* and *LIG4* loci.

#### ENO1-SI

pV1210 containing the gRNA sequence for targeting *ADE2* was digested with KpnI and SacI. The linear CRISPR cassette was purified from a 1% agarose gel using the Wizard® PCR and SV Gel Clean-Up System (Promega). The transformation mixture contained 1 µg of the CRISPR cassette and 3 µg donor DNA.

#### HIS-FLP

pADH99Cau was digested with MssI and the linearized cassette was purified from a 1% agarose gel using the Wizard® PCR and SV Gel Clean-Up System (Promega). Universal fragment A and unique fragment B (gRNA introduction) were generated from pADH100Cau and were stitched together into fragment C using Phusion® High-Fidelity DNA Polymerase (New England Biolabs). The transformation mixture comprised 2 µg of the linearized product of pADH99Cau, 2 µg of fragment C and 3 µg donor DNA.

#### LEUpOUT

pCE35 was digested with MssI and the linearized cassette was purified from a 1% agarose gel using the Wizard® PCR and SV Gel Clean-Up System (Promega). Universal fragment A and unique fragment B (gRNA introduction) were generated from pCE27 and were stitched together into fragment C using Phusion® High-Fidelity DNA Polymerase (New England Biolabs). The transformation mixture comprised 2 µg of the linearized product of pADH99Cau, 2 µg of fragment C and 3 µg donor DNA.

#### EPIC

The transformation mixture comprised 5 µg pJMR19 modified to target *ADE2* as described previously and 5 µg donor DNA.

### Editing and targeting verification

To minimize the number of PCRs needed for verifying transformants, we screened colonies for auxotrophy on drop-out media. This medium contained 1.7 g/L yeast nitrogen base without ammonium sulfate, 5 g/L ammonium sulfate, 2% glucose, 2% agar, 200 µg/mL nourseothricin, and 0.79 g of either CSM, CSM-ade, CSM-leu, or CSM-his (referred to in the manuscript as CSM+NTC, CSM-ade+NTC, CSM-leu+NTC and CSM-his+NTC respectively). Using velveteen replica plating, transformation plates were replicated onto both complete synthetic media and drop-out media lacking specific nutrients: adenine (for all systems), histidine (for HIS-FLP), or leucine (for LEUpOUT). Colonies were counted on both original and replica plates. If the 1:1 dilution plate was overgrown, colony counts from the 1:10 dilution plate were used. Colonies failing to grow on drop-out medium without adenine were processed further with PCR to confirm correct integration. The same approach was used for HIS-FLP and LEUpOUT transformants, screening them for correct cassette integration via PCR. For *ENO1-SI* transformants, colonies from each plate were pooled, and PCR was conducted en masse to confirm the presence of the cassette at the correct locus. Editing of two bases in *ADE2* was confirmed by allele-specific PCR (AS-PCR), following the method described by Carolus *et al.* [56]. The gradient PCR for the selection of annealing temperature is shown in **Figure S7**. AS-PCR was performed with live cells as template and an initial denaturation step at 95°C for four minutes, and 30 cycles of DNA amplification following the standard protocol of Taq DNA polymerase (New England Biolabs). Only transformants that showed a band at the expected molecular weight for the mutant allele and not the wild-type allele were considered correctly edited. Targeting verification of the HIS-FLP and LEUpOUT systems was done by PCR with primer pairs that were designed to bind upstream and downstream of the homologous regions and in the original ORF sequence or the constructed cassette.

## Acknowledgments

This work is part of the CycleDrug project, which has been supported by the Fund for Scientific Research Flanders (FWO) under the framework of the JPIAMR – Joint Programming Initiative on Antimicrobial Resistance fund (G0L1622N), granted to P.V.D. Additionally, this work was supported by a C3 grant from the Industrial Research Fund of KU Leuven (C3/22/007) granted to P.V.D. Researchers H.C. and D.S. were supported by FWO PhD fellowships 11D7620N and 11J8122N respectively. H.C. was also supported by a post-doctoral fellowship granted by KU Leuven Internal Funds (PDMT2/23/032). C.J.N. acknowledges support from the National Institutes of Health (NIH) National Institute of General Medical Sciences (NIGMS) award R35GM124594, and from the Kamangar family in the form of an endowed chair to C.J.N. Researcher C.L.E. was supported by fellowship F31DE028488 from the NIH National Institute of Dental & Craniofacial Research (NIDCR).

## Author contributions

D.S. led the study, carried out experiments and data analysis. H.C. conceptualized and supervised the study, provided funding, carried out experiments and wrote the original draft. A.S. provided experimental advice. C.L.R. carried out experiments. C.L.E., A.D.H., and C.J.N. provided the Clade I.2 wild type strain, constructed the *lig4Δ* strains, and provided experimental advice. J.M.R. constructed and provided the EPIC plasmid and experimental advice. P.V.D. provided funding and supervised the study. H.C. and D.S. contributed equally to the study. All authors contributed to editing the manuscript.

---

**Figure S1:**
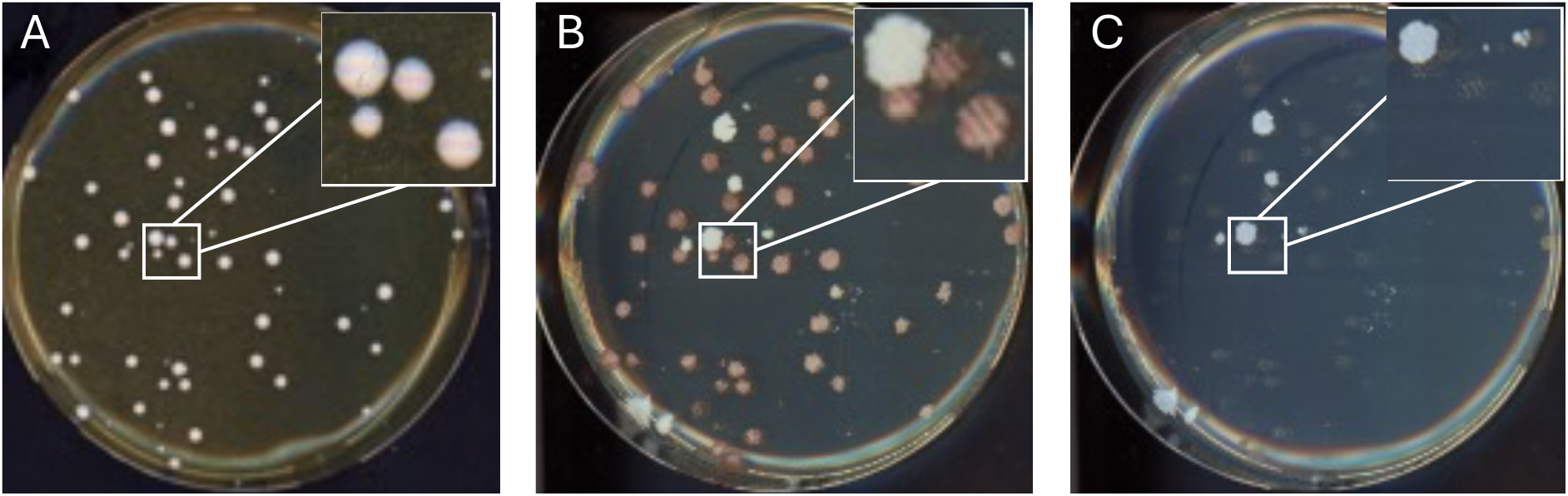
*ADE2* loss of function phenotype. The transformation plate (A) and its replicates in complete synthetic medium (CSM) (B) and dropout-medium lacking adenine (C). Transformants on YPD agar do not show the characteristic red color described in other species. In CSM with minimal amounts of adenine (10 mg/L), the *ade2* transformants develop a brown hue, while in adenine lacking drop-out medium *ade2* transformants are unable to grow.

**Figure S2:**
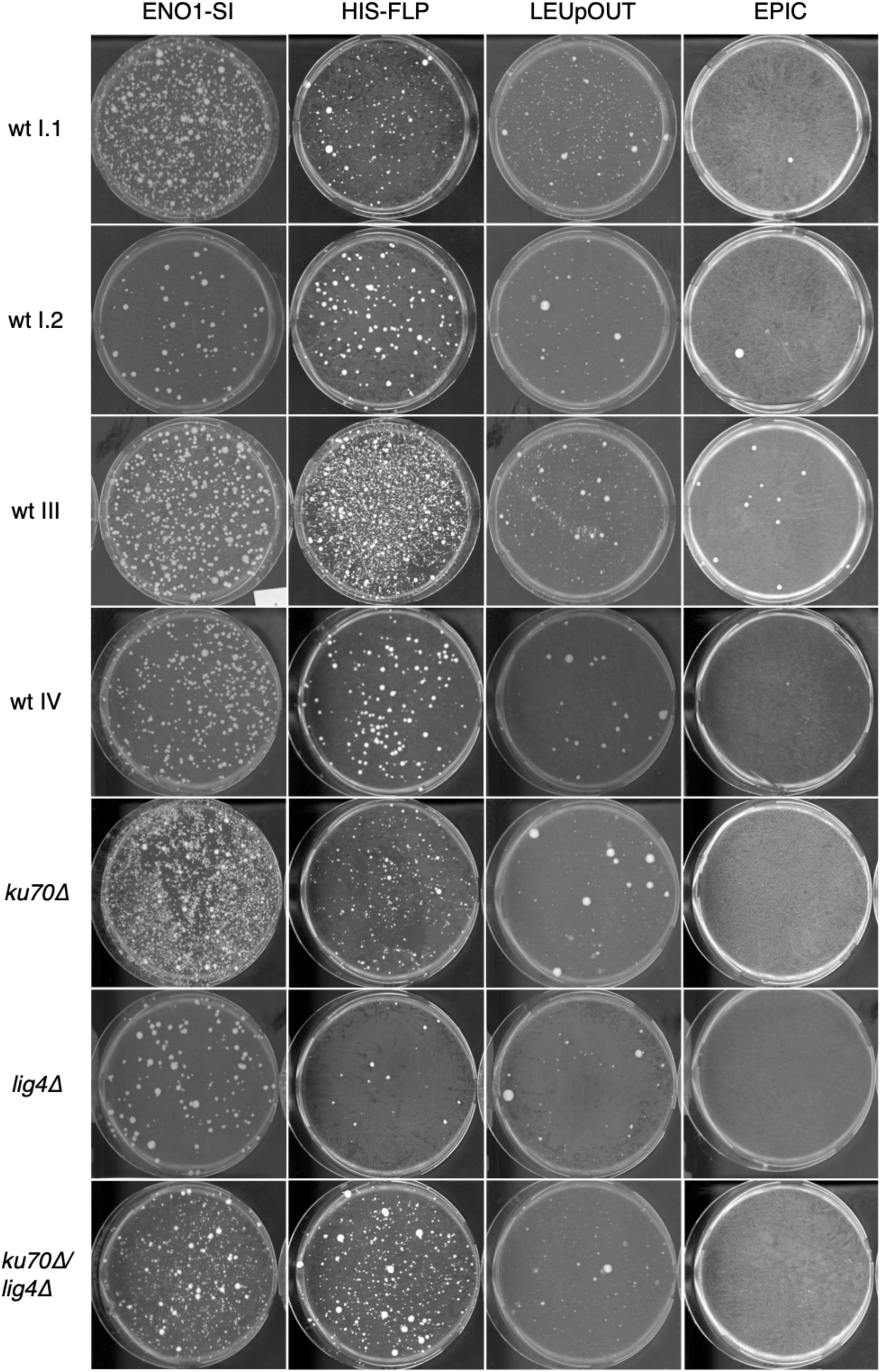
Examples of transformation plates. Representative images of transformation plates (YPD agar with nourseothricin 200 µg/mL) after 2 days incubation at 37°C. One out of three plates for each strain and each system is shown. Microcolonies or background growth was present at variable rates in all integration-based systems and for all strains. Such colonies were excluded from our analysis and further processing.

**Figure S3:**
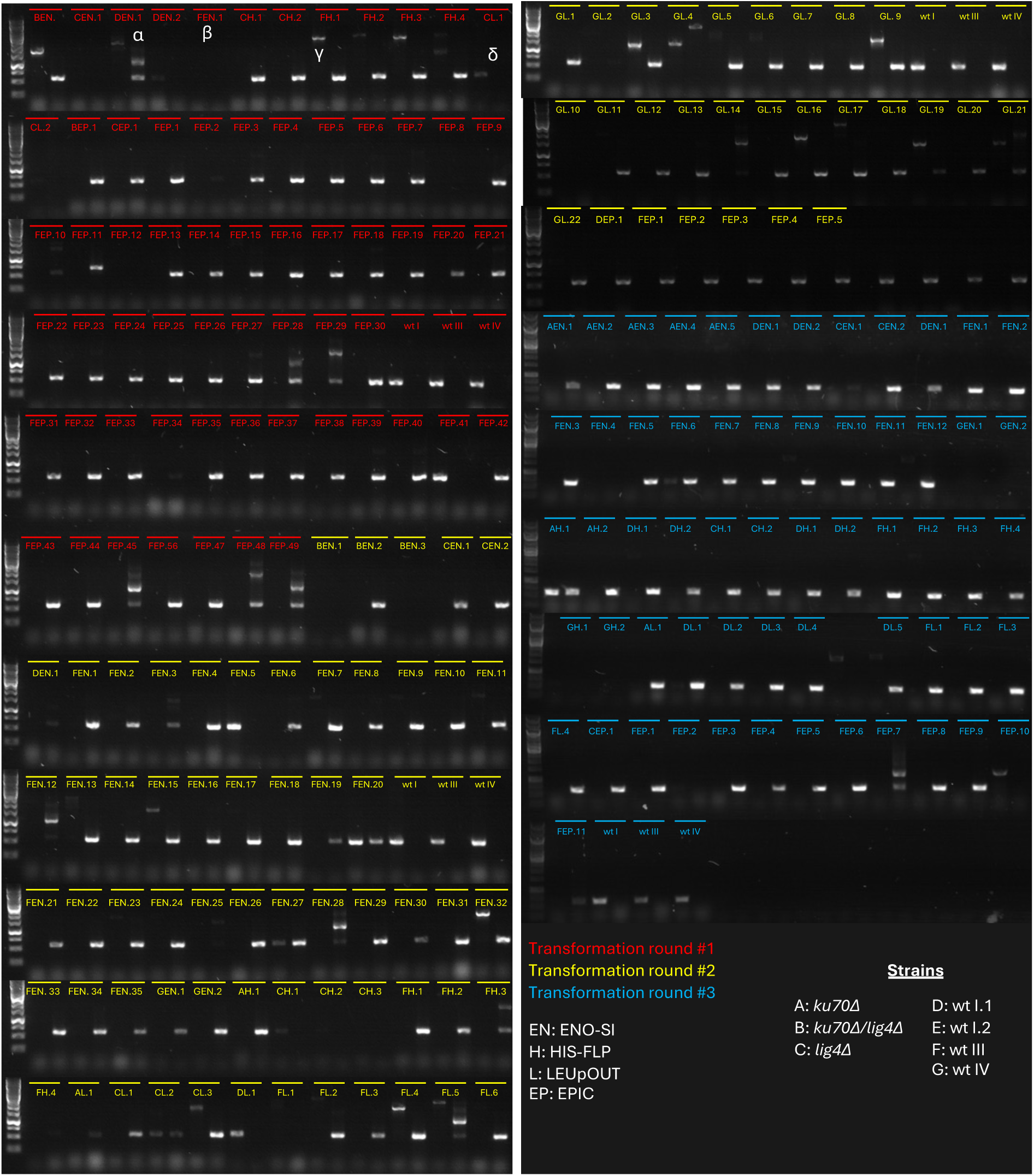
Allele-specific PCRs results for *ADE2* editing efficiency. Agarose gel electrophoresis images showing the PCR products of the auxotrophy-based verified transformants. For each transformant, the wild-type (wt) allele PCR product is loaded in the left lane, followed immediately by the corresponding mutant allele PCR product in the adjacent right lane. The three wt strains were included in every PCR round to ensure the specificity of the primer pairs. Transformants are grouped by transformation round and color-coded, with each sample labelled according to the strain code (A to G), gene editing system code (EN, H, L, or EP), and transformant number. Only transformants with a single band at the expected molecular weight (563 bp) for the mutant allele and no band for the wt allele were considered successfully edited. Transformants showing multiple bands (e.g., α), no bands (e.g., β), high-molecular-weight bands in the wt PCR lane (e.g., γ), or only a wt band (e.g., δ) were deemed incorrectly edited. The GeneRuler 1 kb DNA Ladder (Thermo) is loaded in the leftmost lane of each row.

**Figure S4:**
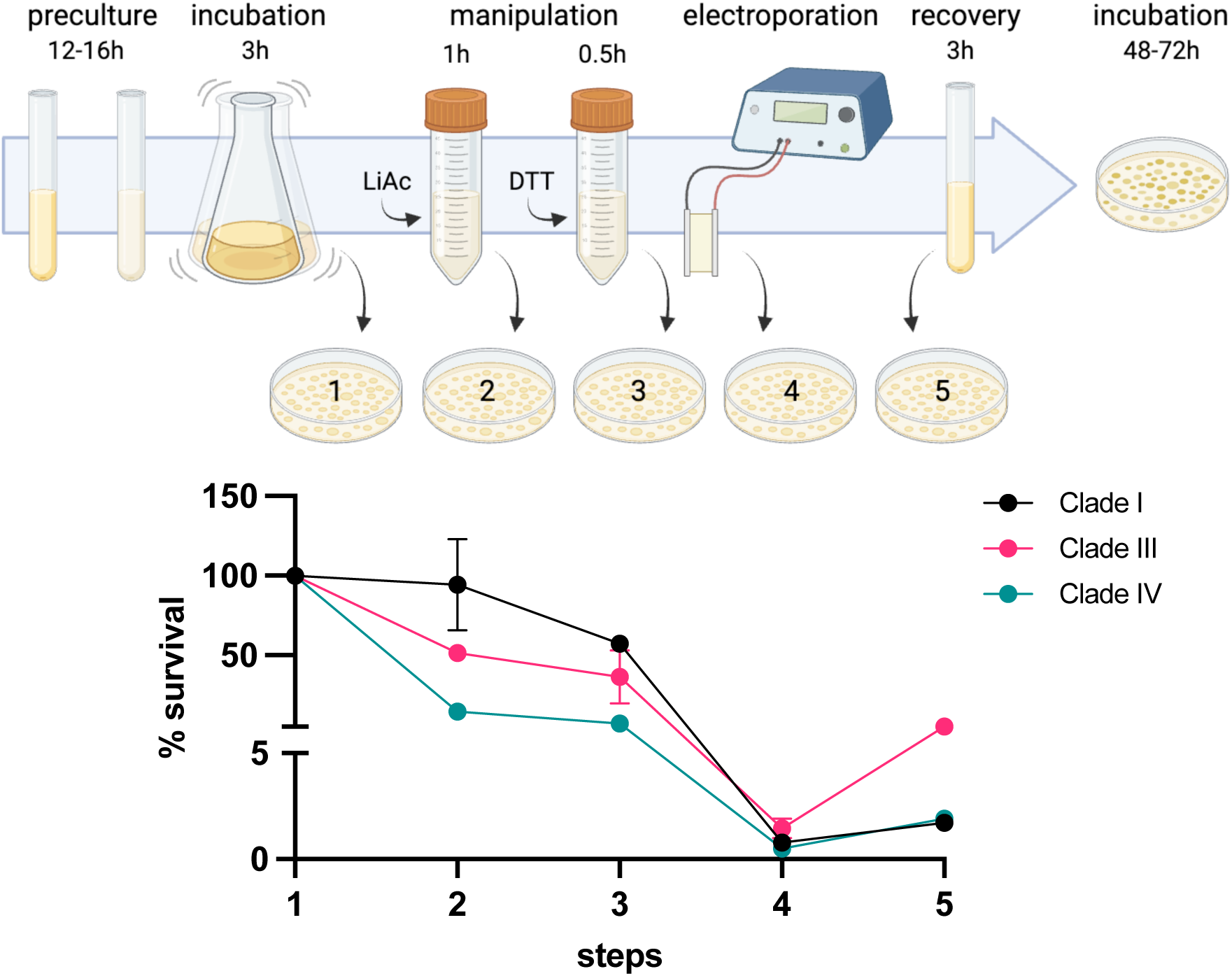
Survival of the wt strains during each transformation step. Relative survival of each wt strain (wt I.1, wt III, and wt IV) after each step of transformation by electroporation. Transformation steps include: (1) three hour incubation in YPD at 37°C, (2) one hour incubation in transformation buffer containing 100 mM lithium acetate at 37°C, (3) thirty minute incubation with 25 mM dithiothreitol (DTT), (4) electric pulse, and (5) three hour recovery in YPD. Data points represent CFU counts from serial dilutions, with survival percentages averaged over two dilutions (10-fold apart). Error bars indicate the standard error of the mean (SEM).

**Figure S5:**
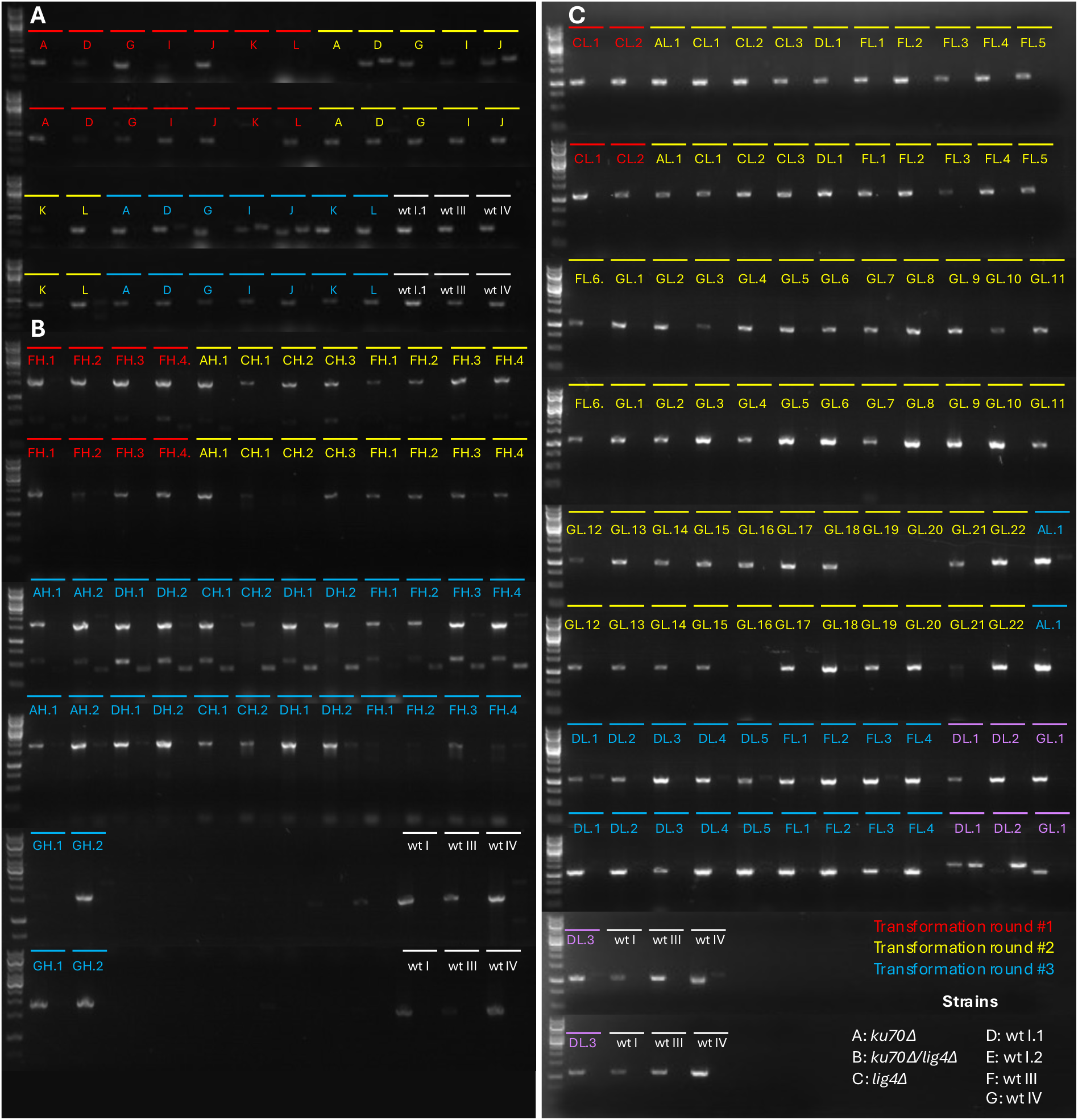
PCR results for cassette integration efficiency. Agarose gel electrophoresis images showing the PCR products of the adenine auxotrophy-based verified transformants. For each transformant, the upstream junction PCR products are loaded in the upper row, while the downstream junction PCR products are loaded in the lower row. In each row, for each transformant, the product of the primer pair binding in the ORF sequence is loaded first, followed immediately by the PCR product of a primer pair binding in the cassette in the adjacent right lane. A) Integration efficiency of the *ENO1-SI* system. A pooled sample of all transformants was used as template DNA for these PCRs. B) Integration efficiency of the *HIS-FLP* system. C) Integration efficiency of the *LEUpOUT* system. Transformants are grouped by transformation round and color-coded, with each sample labelled according to the strain code (A to G), gene editing system code (EN, H, L, or EP), and transformant number. The GeneRuler 1 kb DNA Ladder (Thermo) is loaded in the leftmost lane of each row.

**Figure S6:**
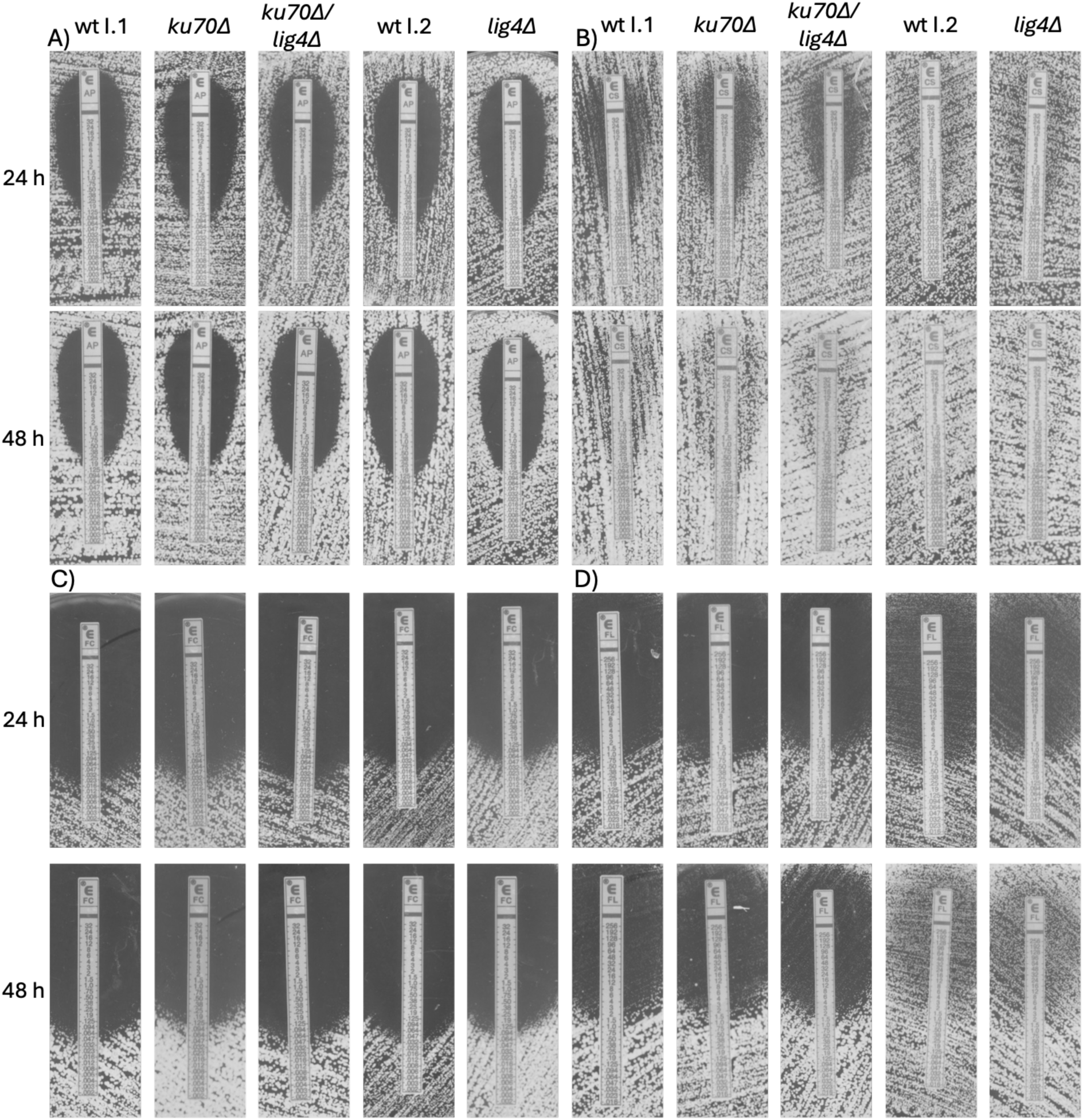
ETEST images for the *ku70Δ*, *ku70Δ/lig4Δ* and *lig4Δ* mutant and their parental strains. ETESTs for amphotericin B (A), caspofungin (B), 5-fluorocytosine (C), and fluconazole (D) are shown. Pictures were taken after 24 and 48 hours incubation at 37 °C. The mutant strains behave similarly to their parental strains. However, the wt I.2 strain and its derivative *lig4Δ* display increased tolerance to caspofungin and fluconazole compared to wt I.2 and its derivatives. Biological replicates of each mutant strain produced consistent results; thus, only one representative replicate is shown.

**Figure S7:**
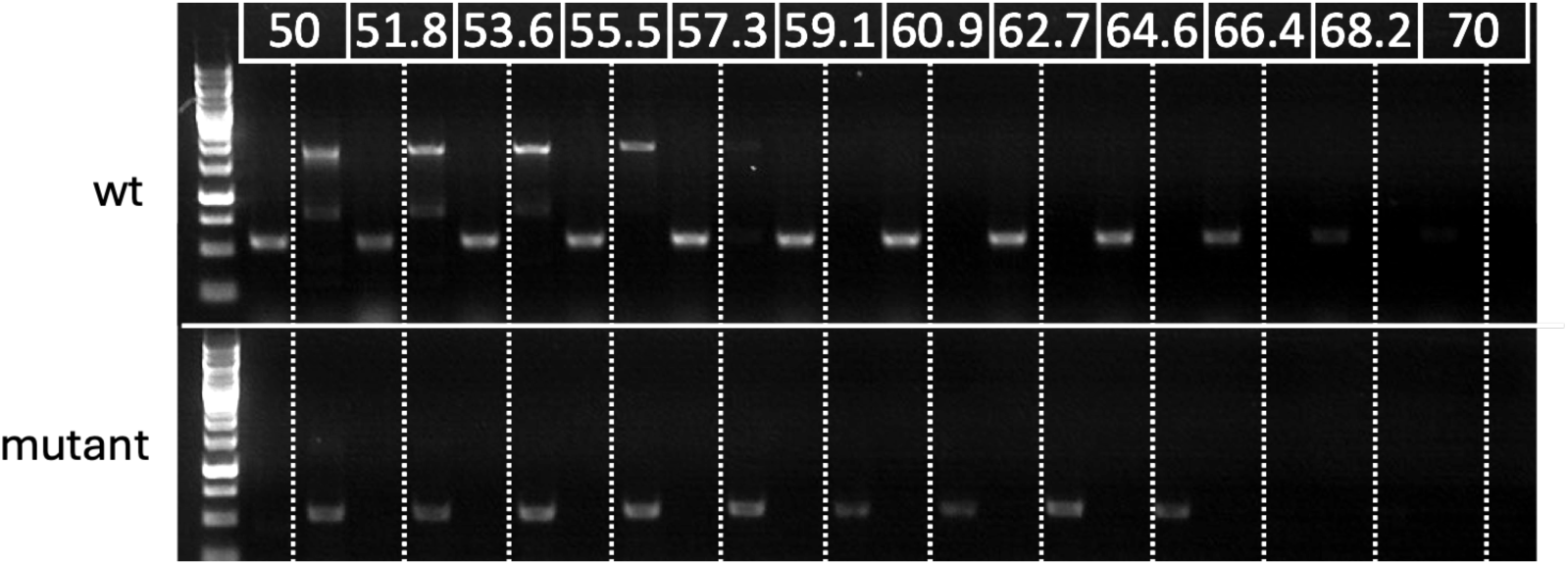
Gradient PCR optimization for Allele-Specific (AS-PCR). Agarose gel electrophoresis images showing the PCR products using a wt strain (wt I.1) and a correct mutant (sequencing verified) using allele-specific primers. Each strain was tested with two primer pairs: one specific to the wild-type (wt) sequence and the other to the mutant sequence. For each strain, the PCR product for the wt allele is shown in the left lane, followed by the mutant allele product in the adjacent right lane. Annealing temperatures used in the thermocycler are indicated above the gel lanes. An annealing temperature of 62.7°C was chosen for subsequent verification of transformants. The GeneRuler 1 kb DNA Ladder (Thermo) is loaded in the leftmost lane of each row.

**Figure S8.**
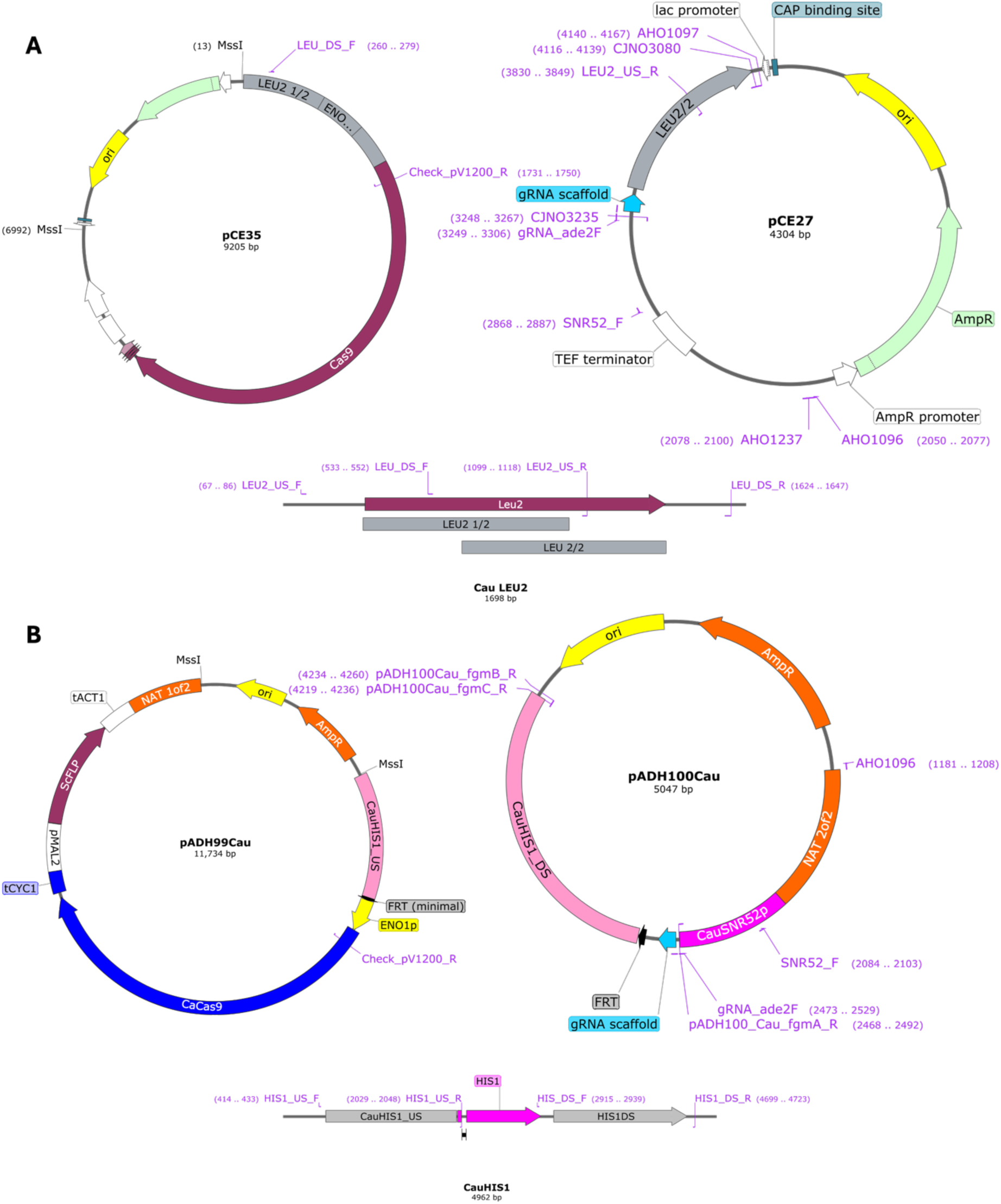
Maps of the plasmids used for *LEUpOUT* (A) and *HIS-FLP* (B). Crucial elements of each CRISPR system are highlighted. Beneath the plasmid maps of each CRISPR system, the genomic locus of *CauLEU2* (A) and *HIS-FLP* (B) are depicted. All primers used to produce the CRISPR linear cassettes, as well as primers used to check the integration of each system are shown.

**Table S1:**
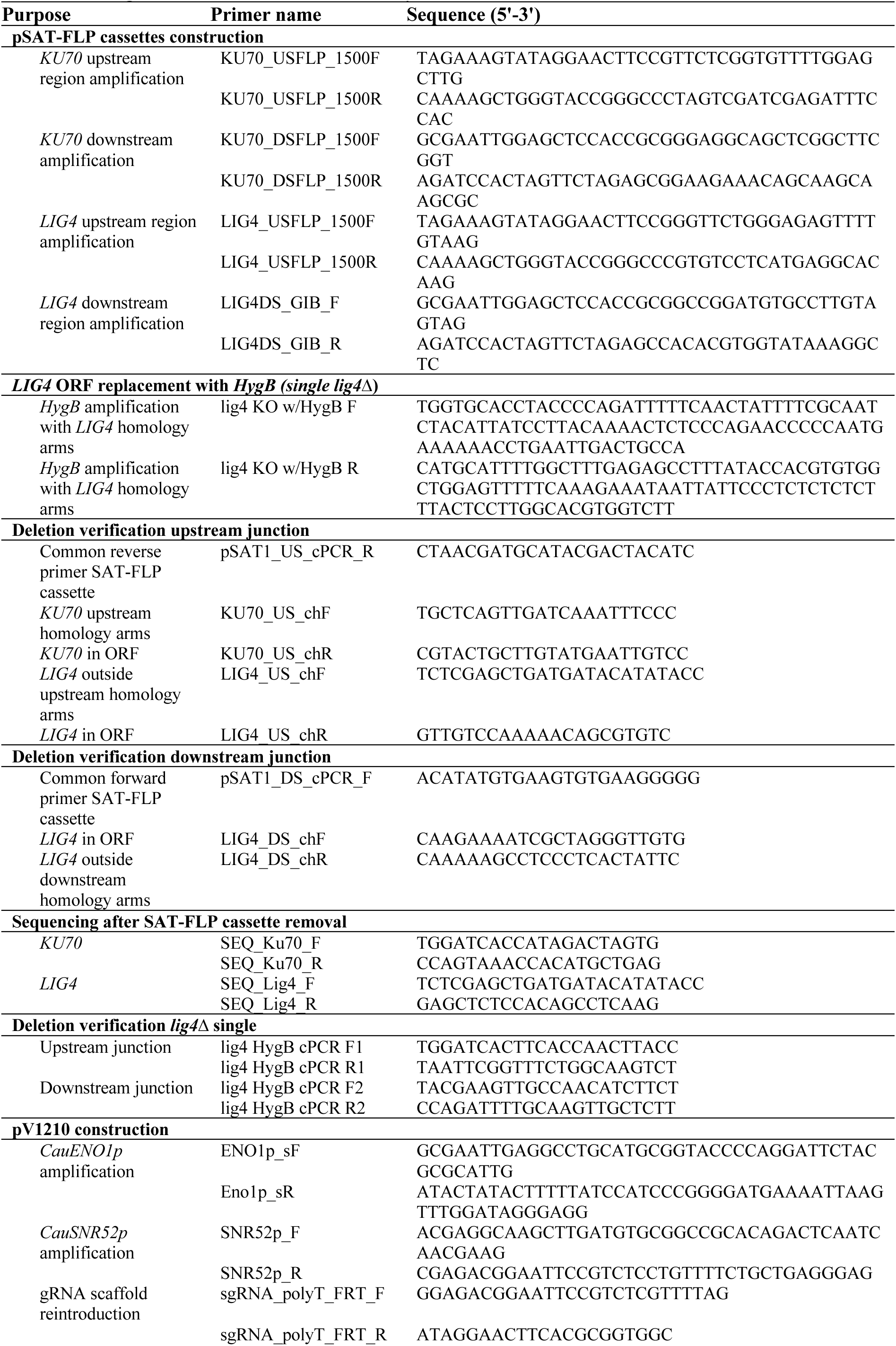
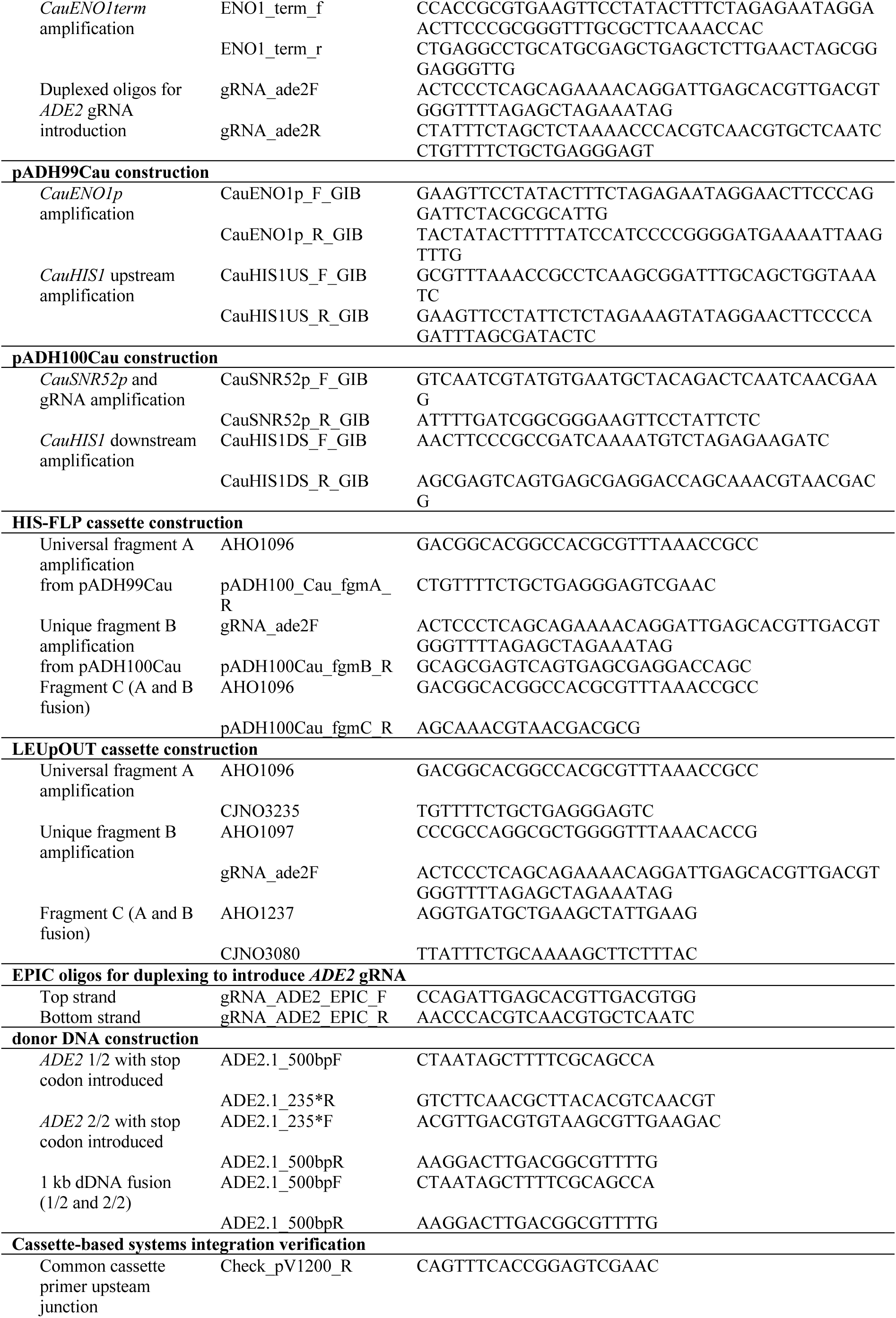
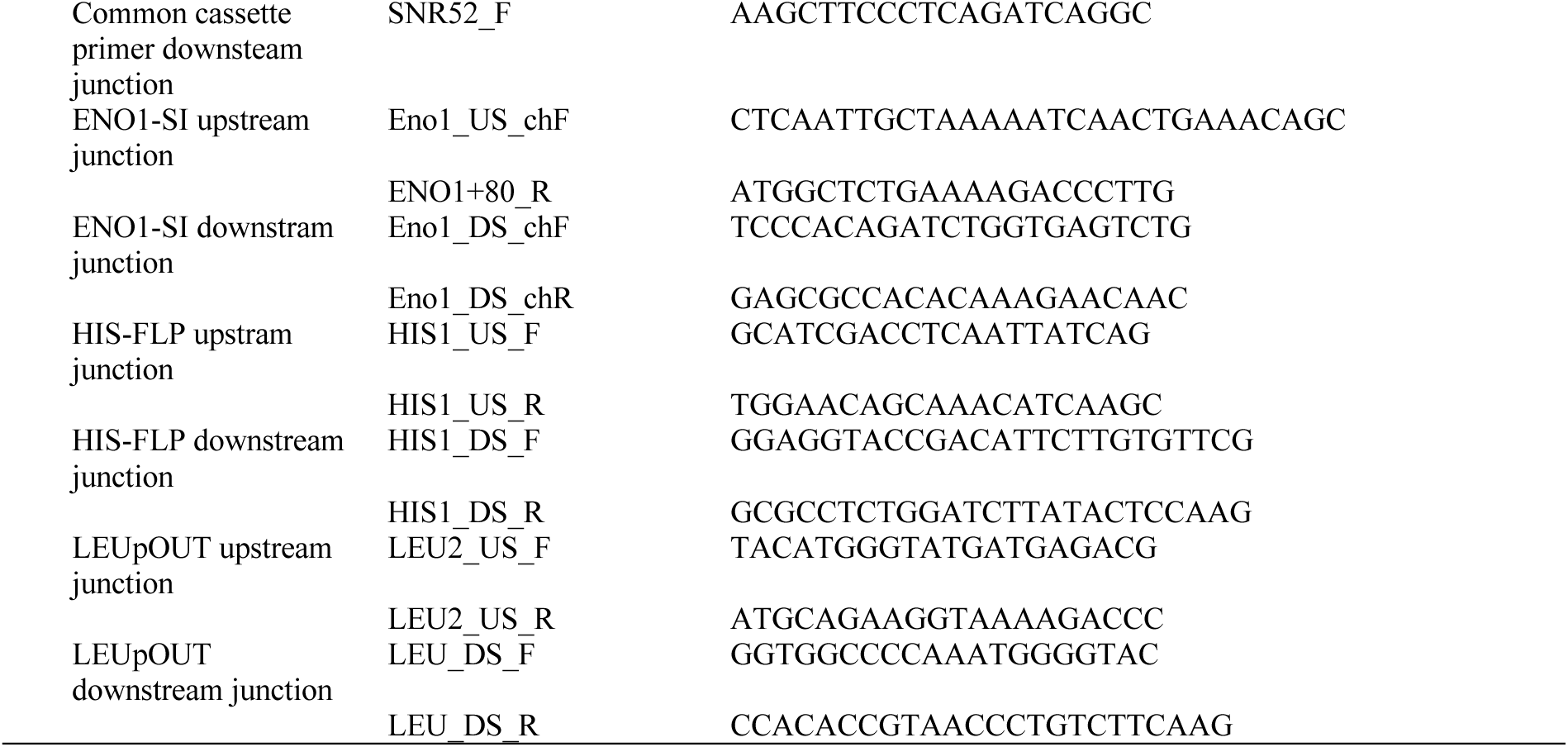
Oligonucleotides used in this study.

**Table S2:**
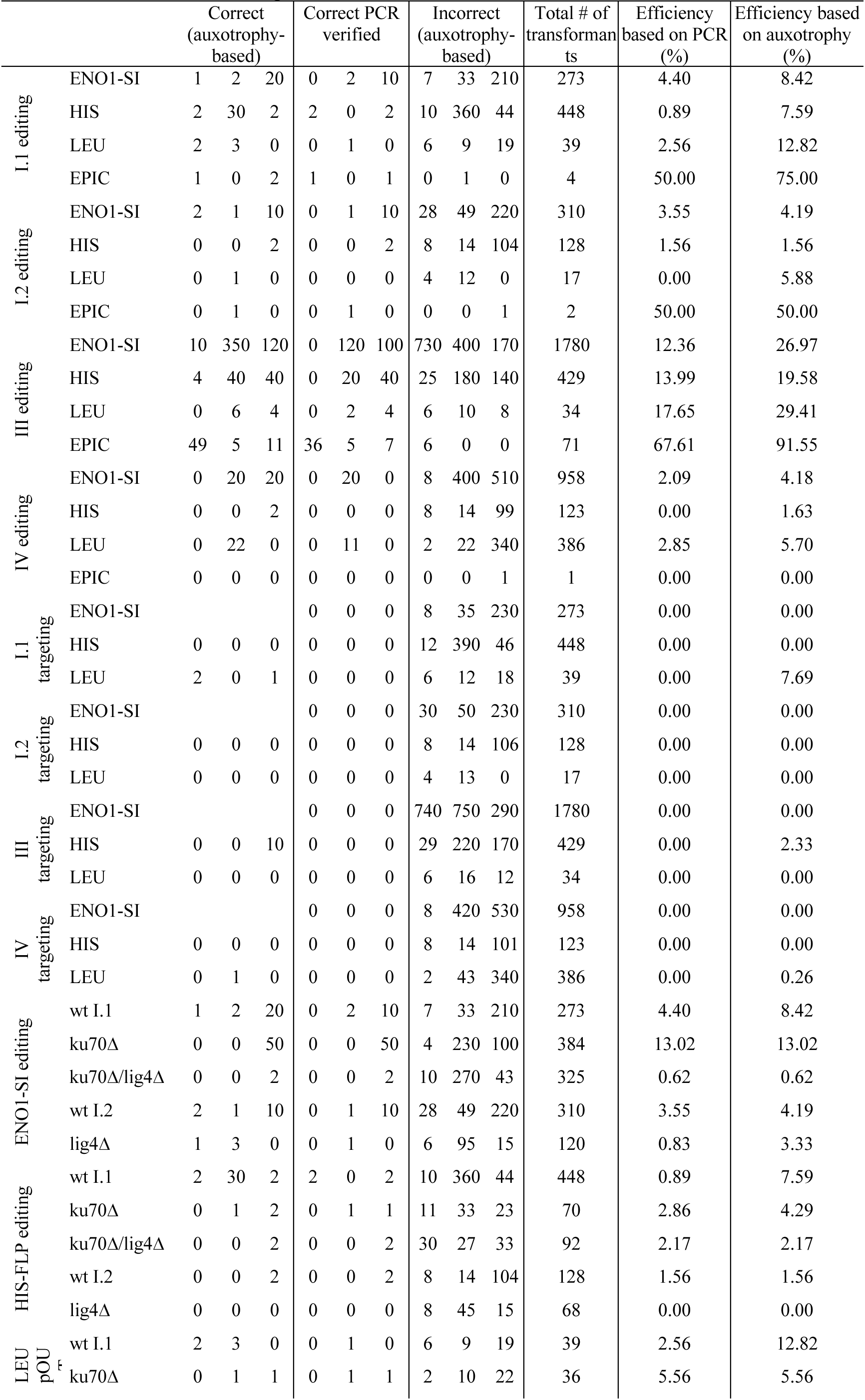
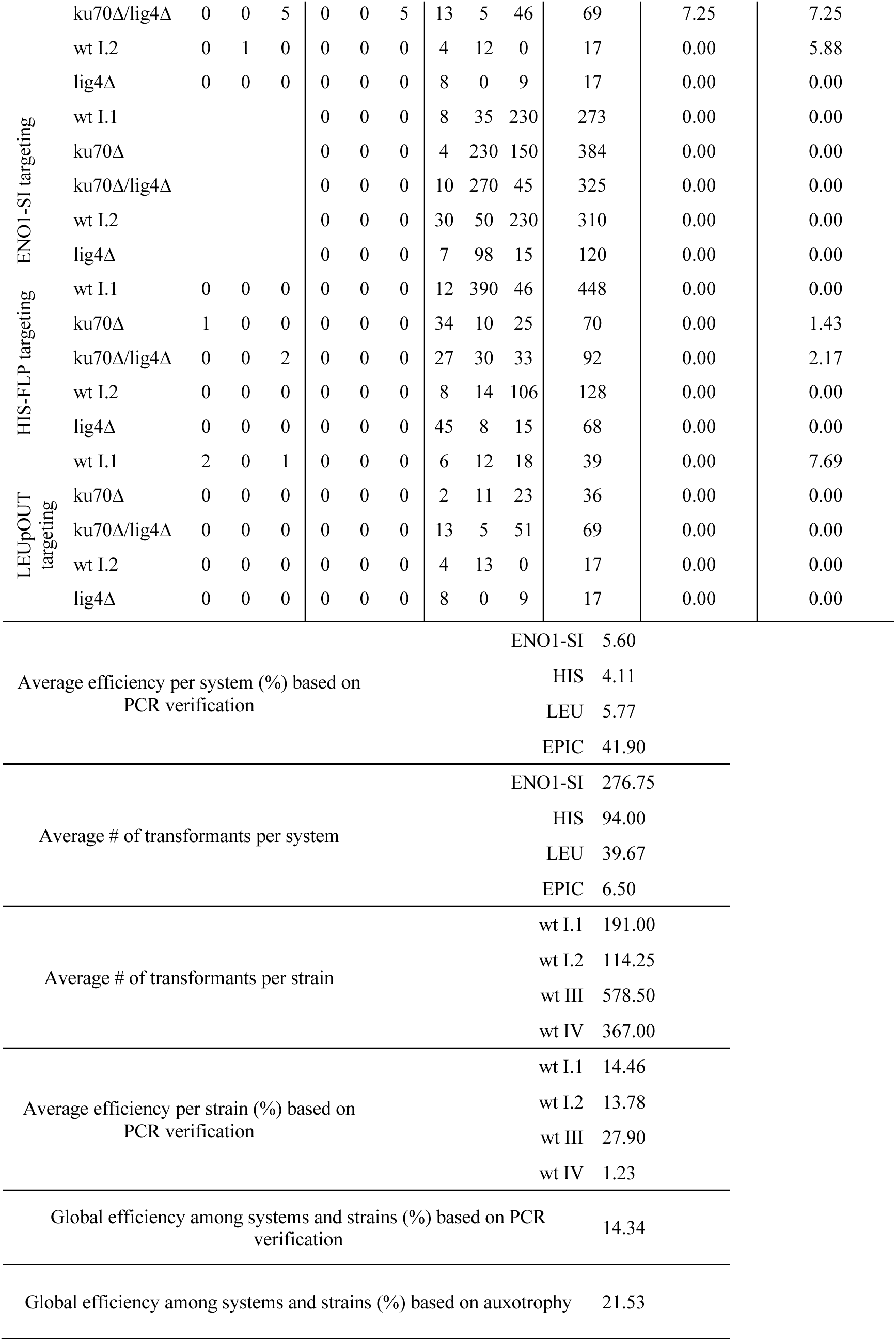
Source data for figures 2, 3 and 5.

**Table S3:** Strains used in this study.

| Strain | strain ID (AR-ID)<sup>ref</sup> | Genetic background/genotype |
| --- | --- | --- |
| wt I.1 | B8441 (AR0387)<sup>[1]</sup> |  |
| wt I.2 | B8441 (AR0387)<sup>[1]</sup> |  |
| wt III | B11223<sup>[2]</sup> |  |
| wt IV | C52710-20 |  |
| *ku70Δ* | DSC001 | B8441 (wt I.1), *ku70Δ* |
| *ku70Δ* | DSC002 | B8441 (wt I.1), *ku70Δ* |
| *ku70Δ* | DSC003 | B8441 (wt I.1), *ku70Δ* |
| *ku70Δ/lig4Δ* | DSC004 | B8441 (wt I.1), *ku70Δ lig4Δ* |
| *ku70Δ/lig4Δ* | DSC005 | B844 (wt I.1), *ku70Δ lig4Δ* |
| *ku70Δ/lig4Δ* | DSC006 | B8441 (wt I.1), *ku70Δ lig4Δ* |
| *lig4Δ* | CEC037 | B8441 (wt I.2), *lig4Δ::HygB* |
| *lig4Δ* | CEC038 | B8441 (wt I.2), *lig4Δ::HygB* |

1. Lockhart SR, Etienne KA, Vallabhaneni S, Farooqi J, Chowdhary A, Govender NP, et al. Simultaneous emergence of multidrug-resistant *Candida auris* on 3 continents confirmed by whole-genome sequencing and epidemiological analyses. *Clinical Infectious Diseases* 2017;64(2):134-40. Epub 2016/12/19. doi: 10.1093/cid/ciw691. PubMed PMID: 27988485; PubMed Central PMCID: PMC5215215.
2. de Groot T, Puts Y, Berrio I, Chowdhary A, Meis JF, Heitman J. Development of *Candida auris* Short Tandem Repeat Typing and Its Application to a Global Collection of Isolates. *mBio*. 2020;11(1):e02971-19. doi: 10.1128/mBio.02971-19.

